# IFIT3 Associates with m⁶A-Modified RNA to Restrict Hepatitis C Virus Infection

**DOI:** 10.64898/2026.03.25.714224

**Authors:** Matthew Thompson, Moonhee Park, Netanya S. Schlamowitz, Matthew R. Lanahan, Yunsum Nam, Stacy M. Horner

## Abstract

Interferon-induced proteins with tetratricopeptide repeats (IFITs) are RNA-binding effectors that restrict infection by diverse RNA viruses. Among the IFIT family, how IFIT3 recognizes RNA remains the least understood. Here, we identify IFIT3 as preferentially associating with N6-methyladenosine (m⁶A)–modified hepatitis C virus (HCV) genomic RNA and host transcripts to restrict HCV infection. IFIT3 cellular RNA binding sites and m⁶A sites, mapped transcriptome-wide by HyperTRIBE-seq during HCV infection, showed significant overlap. This m⁶A preference was further supported by findings that IFIT3 binding sites significantly overlapped those of established m⁶A-binding proteins; that inhibiting m⁶A installation reduced IFIT3 association with m⁶A-modified HCV RNA and cellular transcripts; that IFIT3 co-purified more efficiently with m⁶A-modified short RNA probes than with unmodified controls; and that mutating m⁶A consensus motifs in the HCV genome reduced IFIT3 association with viral RNA. Structure–function analyses identified two regions required for RNA probe binding: tetratricopeptide repeat domains 1–2 (TPR1–2) and a previously uncharacterized predicted helical hairpin between TPRs 6 and 7. In infected cells, the helical hairpin was required for IFIT3 association with HCV RNA but dispensable for interactions with other IFIT proteins. Conversely, TPR1–2 was dispensable for HCV RNA binding but essential for IFIT2 interaction, establishing that these functions are structurally separable. Loss of either region diminished antiviral activity, as indicated by increased levels of HCV RNA in clarified supernatants. Consistent with models of m⁶A-linked restriction of late stages of infection, extracellular HCV RNA showed reduced m⁶A and decreased IFIT3 association relative to intracellular RNA. Together, these findings define an m⁶A-linked mechanism by which IFIT3 engages viral RNA and reveal an unexpected role for m⁶A in antiviral effector function.

**SIGNIFICANCE:** RNA-binding proteins are critical effectors of antiviral defense, yet for many of these proteins the mechanisms of viral RNA recognition remain unclear. Here, we show that m^6^A modification on both host and viral RNA promotes recognition by the interferon stimulated gene, IFIT3. RNA recognition did not require interaction with IFIT1 or IFIT2, although IFIT3 antiviral function required both RNA binding and interaction with IFIT2. These findings identify m^6^A as a new regulator of IFIT protein function and broaden our understanding of how RNA modifications shape antiviral restriction.

## INTRODUCTION

RNA-binding proteins (RBPs) act as diverse effectors in the cell-intrinsic innate immune response to RNA viruses. The foundation of this defense lies at the host-virus interface, where cells have evolved mechanisms to sense and inhibit viral RNA^1^. RBP-mediated restriction occurs either indirectly, by inducing expression of additional effectors that initiate or amplify signaling, or directly, by targeting specific phases of the viral lifecycle^2–4^. These mechanisms often overlap, such that RNA-sensing RBPs can induce interferons (IFN) that in turn drive expression of IFN-stimulated genes (ISGs), which can encode antiviral RBPs. While the sensing and signaling functions of antiviral RBPs are well defined^1,5^ the mechanisms underlying direct restriction by many antiviral RBPs remain less well understood. Among the ISGs, the IFIT proteins have emerged as RBPs with diverse RNA binding preferences that regulate infection by a wide-array of viruses^6,7^. However, our understanding of IFIT protein function and mechanism remains incomplete.

The IFIT family, among the most highly expressed ISGs^8^, consists of four major human isoforms: IFIT1, IFIT2, IFIT3, and IFIT5^6,7^. These proteins share high sequence conservation and contain several tetratricopeptide repeat (TPR) domains^9,10^. All IFIT proteins have been described as RBPs that regulate viral infection^7,11–16^, yet studies have revealed unique functions for both individual and specific heterocomplexes^12,16,17^. For example, IFIT1 prevents translation initiation by binding directly to the 5’ cap of RNA lacking 2’-*O*-methylation at the first nucleotide (cap0), a mechanism that restricts several flaviviruses^6,7^. In contrast, IFIT2 preferentially binds AU-rich RNA sequences and exhibits both pro- and antiviral roles by either promoting or inhibiting viral translation^14,18,19^. IFIT5 binds both 5’ cap structures and AU-rich RNA, though its ligand affinities and viral restriction profiles differ from other IFITs^13,20–23^. The RNA binding properties and subsequent function of IFIT3 remain the least characterized of the IFIT family. Generally, IFIT3 has been considered a cofactor that enhances the RNA binding and antiviral properties of IFIT1 or IFIT2 through complex formation^16,17^. Consequently, IFIT3 pro- and antiviral functions often mirror those of IFIT1 and IFIT2^7,14^. However, recent work has begun defining a role for IFIT3 directly binding RNA to contribute to IFIT family viral restriction.

Two recent studies have demonstrated IFIT3 binds to viral RNA to regulate infection via different mechanisms. In one example, it was shown that murine IFIT3 directly binds host and viral RNA in a heterodimeric complex with IFIT2 that inhibits translation of viral mRNAs bearing short 5’ UTRs^12^. Conversely, IFIT3 binding to influenza A virus mRNA was shown to enhance viral translation^11^. These new findings suggest an underappreciated role for IFIT3 binding viral RNA and motivate re-examining the role of IFIT3 in previously described viral models. For example, IFIT3, along with IFIT1 and IFIT2 has previously been observed to restrict hepatitis C virus (HCV), but how the IFIT proteins recognized HCV RNA and whether IFIT3 RNA binding is required, was not explored^15,24^. Moreover, it remains unclear what RNA features, such as sequence context, secondary structure, and modification state regulate IFIT3 RNA binding.

Here, we define how IFIT3 engages RNA and contributes to antiviral defense during HCV infection. Using transcriptome-wide mapping, we identified IFIT3 binding sites on host and viral RNAs and found that IFIT3 binding patterns closely resemble those of the RNA binding “reader” proteins the recognize the N⁶-methyladenosine (m^6^A) RNA modification. Consistent with this, inhibiting m^6^A deposition reduced IFIT3 association with both the HCV genome and cellular RNAs, and IFIT3 preferentially co-purified with m^6^A-modified RNA probes. Domain and mutational analyses further showed that IFIT3 RNA binding is mediated by a region independent of IFIT1/IFIT2 interactions. Functionally, both RNA binding and IFIT2 interaction were necessary for maximal restriction of HCV infection, as disrupting either activity reduced antiviral potency. These results identify m^6^A-dependent viral RNA binding by IFIT3 as a new mechanism contributing to HCV restriction.

## RESULTS

### IFIT3 recognizes host and viral RNA with m⁶A reader–like binding preferences

To define IFIT3 RNA binding during HCV infection, we applied HyperTRIBE-seq, a method that detects both direct and indirect RNA-protein interactions through ADAR-mediated RNA editing^25,26^. An IFIT3-ADAR fusion protein containing the full-length human IFIT3 sequence and the catalytic domain of hyperactive ADAR (E488Q)^27^ was stably expressed in Huh7 cells during HCV infection under a low-dose of IFN treatment to induce any potential cofactors (**Figure 1A, Supplementary Figure 1A**). Importantly, these experiments also included a matched control cell line expressing only the catalytic domain of hyperactive ADAR (referred to as ADAR-ctl). ADAR-ctl edits were used to generate a background dataset to control for ADAR-specific RNA binding biases^26,28^. Bulk intracellular RNA from these cells was analyzed by RNA-seq and processed using the Bullseye pipeline^29–31^ to identify transcriptome-wide adenosine-to-inosine (A:I) editing events. Editing events were filtered for sites with ≥8% editing in IFIT3-ADAR cells. In cases where editing was detected in both IFIT3-ADAR and ADAR-ctl cells, filtering required ≥8% editing and ≥1.3 fold enrichment relative to ADAR-ctl. With stringent filtering for edits common between all 3 replicates, this analysis identified 1830 IFIT3-driven editing events across 1271 unique genes (**Figure 1B**). Gene ontology analysis revealed enrichment for membrane-associated compartments, including the Golgi apparatus, lysosomes, and endoplasmic reticulum (**Figure 1C**).

**Figure 1.**
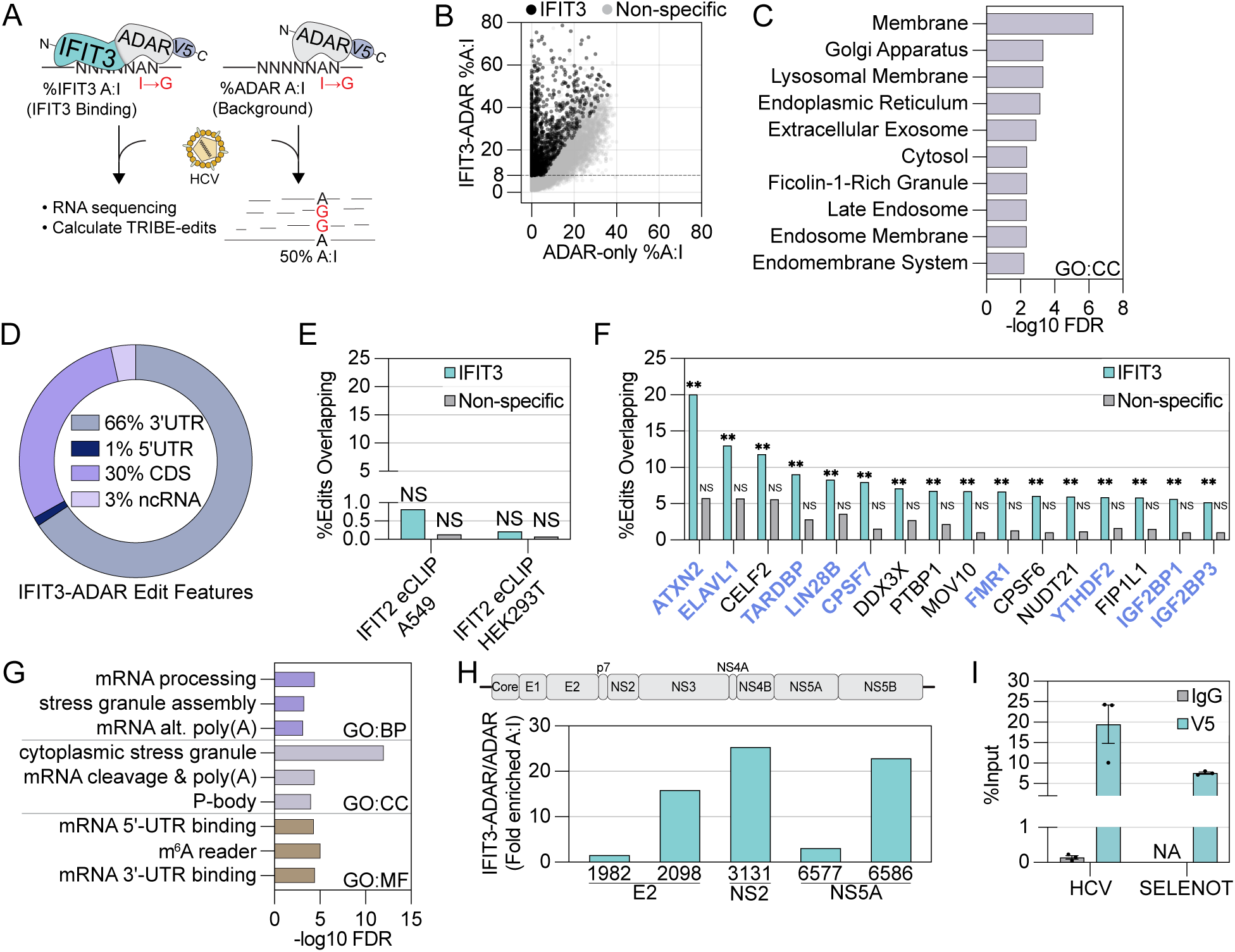
IFIT3 recognizes host and viral RNA with m⁶A reader–like binding preferences. (**A**) Schematic depicting HYPERTRIBE-seq design in which a V5-tagged ADAR catalytic domain is expressed alone or fused to WT IFIT3, followed by RNA sequencing and quantification of ADAR-editing via A to G mutation. (**B**) Comparison of IFIT3-ADAR and ADAR-only %A:I editing at sites with ≥1% editing in IFIT3-ADAR cells. Sites that passed all filtering and statistical cutoffs are indicated in black. (**C**) Top 10 functional enrichment categories for IFIT3-ADAR edit sites. (**D**) Donut chart depicting genomic content of RNA in which IFIT3 edits were detected. (**E**) % of IFIT3-ADAR edits or non-specific edits that fell within published IFIT2 CLIP-seq data from two cell types: 293T^32^ and A549^14^. (**F**) Postar3 CLIPdb datasets in which 5% or more of IFIT3 edit sites fell within a CLIP peak and had an FDR < 0.05 after permutation testing (1,000 shuffles) of the difference between IFIT3 and Non-specific edit site overlap. (**G**) Functional enrichment categories for overlapping proteins in (F). (**H**) Fold enrichment of %A:I edits on HCV RNA that passed all filtering and statistical cutoffs in IFIT3-ADAR cells compared to ADAR-only cells. (**I**) Native RIP-qPCR of IFIT3-V5 or IgG control from HCV-infected Huh7 (72 hpi, MOI 0.3) cells stably expressing IFIT3-V5. Values represent % of total RNA pulled down by V5 antibody or IgG. See also Figure S1.

**Supplementary Figure 1.**
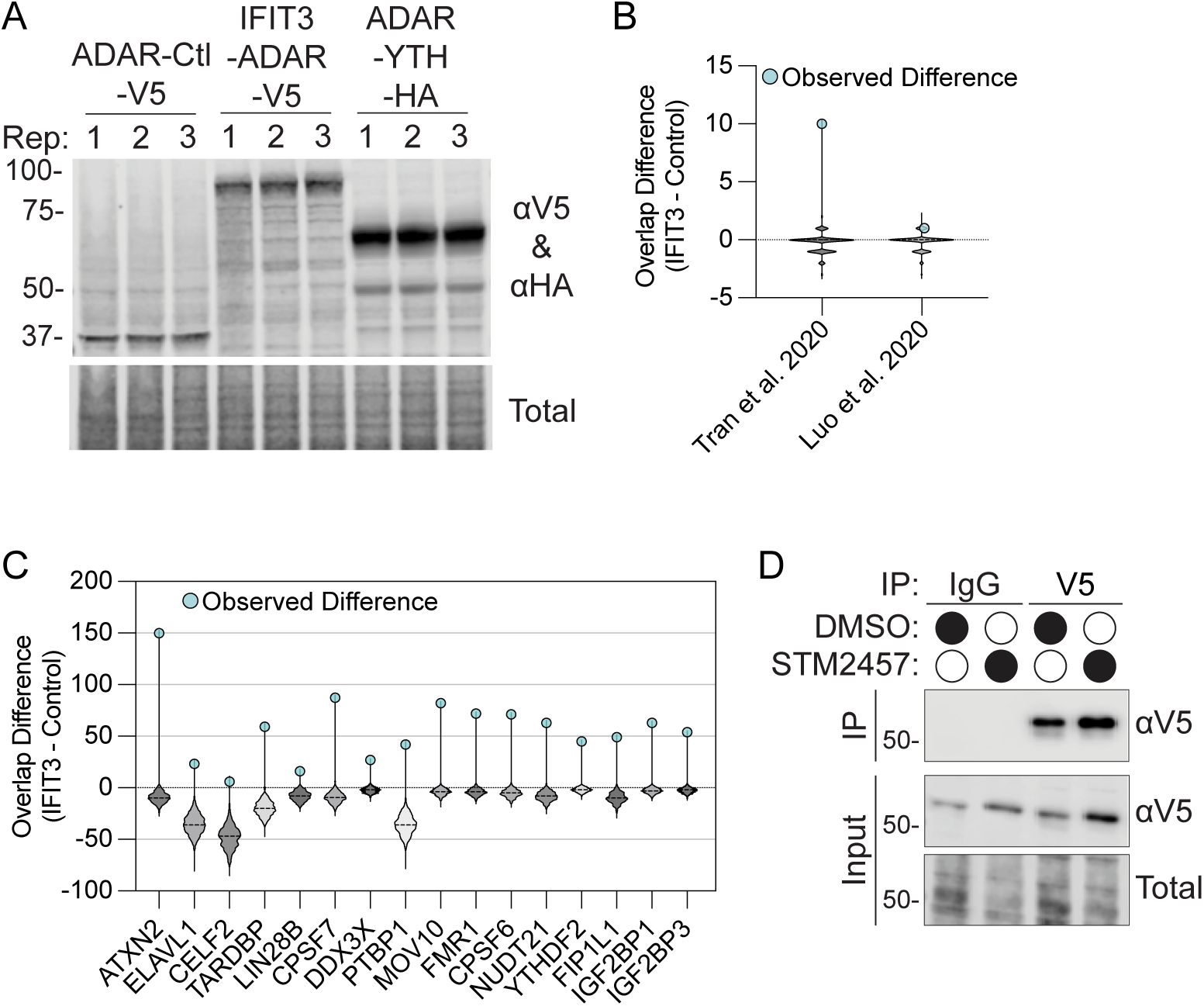
Validation of TRIBE-seq fusion constructs and supporting overlap analyses, related to Figures 1 and 2. **(A)** Immunoblotting of stably expressed ADAR-Ctl-V5, IFIT3-ADAR-V5, and ADAR-YTH-HA fusion proteins in Huh7 cells used for TRIBE-seq experiments. **(B)** Permutation test distributions (1,000 shuffles) for overlap of IFIT3-ADAR–specific or non-specific edit sites with published IFIT2 eCLIP datasets from HEK293T (Luo et al., 2020) and A549 (Tran et al., 2020) cells, with observed overlap difference (IFIT3 − Control) indicated. **(C)** Permutation test distributions (1,000 shuffles) for overlap of IFIT3-ADAR–specific or non-specific edit sites with indicated POSTAR3 CLIPdb datasets. The observed overlap difference (IFIT3 − Control) is shown for each RNA-binding protein. **(D)** Immunoblotting of IFIT3-V5 expression and immunoprecipitation from HCV-infected Huh7 cells stably expressing IFIT3-V5 treated with DMSO or 20 μM STM2457, corresponding to RIP-qPCR experiments in in Figure 1I and Figure 2A.

To assess IFIT3 binding preferences, we annotated genomic features containing IFIT3-ADAR editing sites. These sites are located predominantly within 3’ untranslated regions (3′UTR) and coding sequences (CDS) (**Figure 1D**). Because IFIT2 has been described to show a similar binding pattern, we compared the IFIT3-ADAR dataset to previously published IFIT2 eCLIP data from other cell types^14,32^, and used a windowed overlap empirical permutation test (100 bp window, 1,000 shuffles, FDR<0.05) to assess overlap significance. However, we observed no significant overlap between target genes (**Figure 1E, Supplementary Figure 1B).** Nonetheless, IFIT3-ADAR editing sites have significant overlap with previously published CLIP-seq datasets for other RNA-binding proteins^33^, as assessed using permutation tests (**Figure 1F and Supplementary Figure S1C**). These included proteins with defined 3’UTR or m^6^A-associated binding preferences, such as cleavage and polyadenylation factors (CPSF6, CPSF7, FIP1L1, NUDT21)^34^ and canonical m^6^A-associated reader proteins (YTHDF2, IGF2BP2, IGF2BP3)^35,36^ (**Figure 1F-G**). IFIT3 binding sites also overlapped with several non-canonical m⁶A-associated reader proteins, including ELAVL1 (HuR)^37^, TARDBP (TDP-43)^38^, MOV10^39^, and FMR1^40,41^ (**Figure 1F**).

To determine whether IFIT3 also bound viral RNA, we mapped IFIT3-ADAR editing events to the HCV genome, which contains m^6^A^42,43^. We identified five editing sites within the viral coding region corresponding to the E2, NS2, and NS5A genes (**Figure 1H**). Consistent with these results, RNA immunoprecipitation followed by qPCR (RIP-qPCR) of stably expressed IFIT3-V5 (without ADAR fusion) in HCV-infected Huh7 cells revealed enrichment for both HCV RNA and the host m^6^A-modified transcript *SELENOT* **(Figure 1I, Supplementary Figure 1D**). Together, these data demonstrate that IFIT3 associates with broad range of host RNAs with enrichment in 3’UTRs and overlaps with m^6^A-reader-associated regions, as well as with HCV RNA.

### IFIT3 associates with m^6^A-modified RNA transcriptome-wide

Given IFIT3-bound RNAs are also targeted by m^6^A readers, we asked whether its RNA association is m⁶A-dependent. We assayed this using RIP-qPCR following treatment of HCV-infected Huh7 cells with the METTL3 inhibitor STM2457^44^ or DMSO control (**Figure 2A**). IFIT3 associated with HCV RNA, the highly IFIT3-ADAR edited host transcript *SELENOT*, and non-m^6^A-modified transcript *HPRT1*^45^. However, METTL3 inhibition reduced IFIT3 pulldown of HCV RNA and *SELENOT*, but not *HPRT1*, indicating that IFIT3 association with these RNAs depends on m^6^A. To independently assess m^6^A-binding preferences transcriptome-wide, we again applied HyperTRIBE-seq using the previously described ADAR-YTH fusion protein to map m⁶A sites transcriptome wide (**Supplementary Figure 1A**)^29^. Using the same Bullseye pipeline, filtering criteria, and background ADAR-ctl dataset as for IFIT3-ADAR, we identified 3,315 ADAR-YTH edits across 2082 genes (**Figure 2B, Table 1**). When compared with our previously generated m^6^A maps in Huh7 cells^46^ and CLIPdb^33^ datasets, ADAR-YTH edits have significant overlap with m^6^A-methylated RNA immunoprecipitation and sequencing (meRIP-seq) peaks and showed enrichment among m^6^A-reader CLIP-seq sites (100 bp window, 1,000 shuffles, FDR<0.005) (**Figure 2C, Supplementary Figure 2A**).

**Figure 2.**
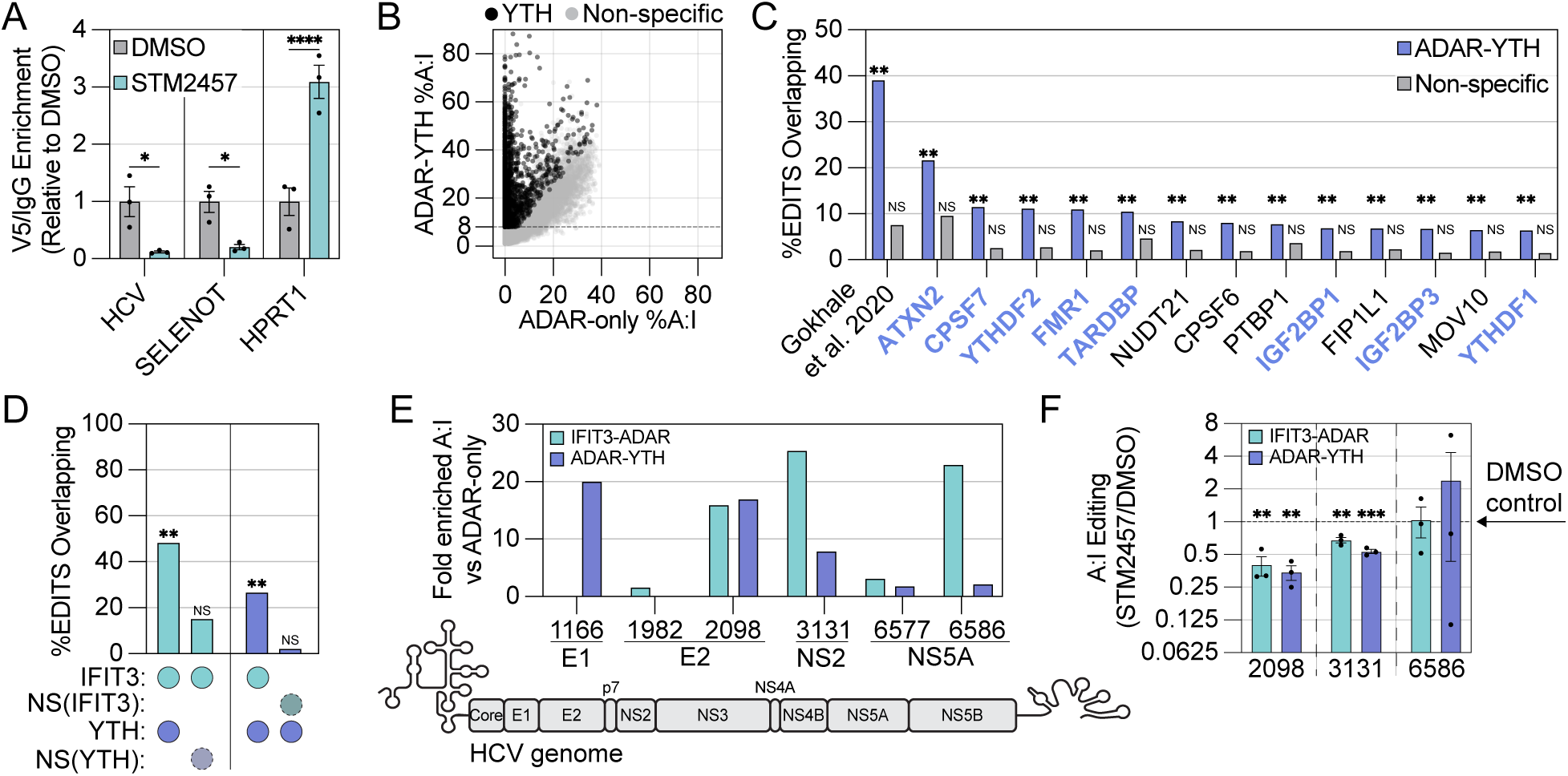
IFIT3 associates with m^6^A-modified RNA transcriptome-wide. (**A**) Native RIP-qPCR of IFIT3-V5 or IgG control from HCV-infected (72 hpi, MOI 0.3) Huh7 cells stably expressing IFIT3-V5 and treated with DMSO or 20 µM STM2457. Values represent enrichment of % total RNA pulled down by V5 antibody compared to IgG followed by normalization to DMSO condition. (**B**) Comparison of YTH-ADAR and ADAR-only %A:I editing at sites with ≥1% editing in IFIT3-ADAR cells. Sites that passed all filtering and statistical cutoffs are indicated in black. (**C**) Postar3 CLIPdb datasets in which 5% or more of ADAR-YTH edit sites fell within a CLIP peak and had an FDR < 0.05 after permutation testing (1,000 shuffles) of the difference between ADAR-YTH and Non-specific edit site overlap. (**D**) Overlap analyses of specific and non-specific edit sites derived from IFIT3-ADAR and ADAR-YTH. Statistical testing was performed via permutation test (1,000 shuffles) with FDR adjustment. (**E**) Fold enrichment of %A:I edits on HCV RNA that passed all filtering and statistical cutoffs in IFIT3-ADAR or ADAR-YTH cells compared to ADAR-only cells. (**F**) RT-PCR Sanger sequencing of HCV TRIBE sites in the context of STM2457 or DMSO treatment. Values are representative of A:I editing in STM2457 treated samples normalized to DMSO treated samples. See also Figures S2.

**Supplementary Figure 2.**
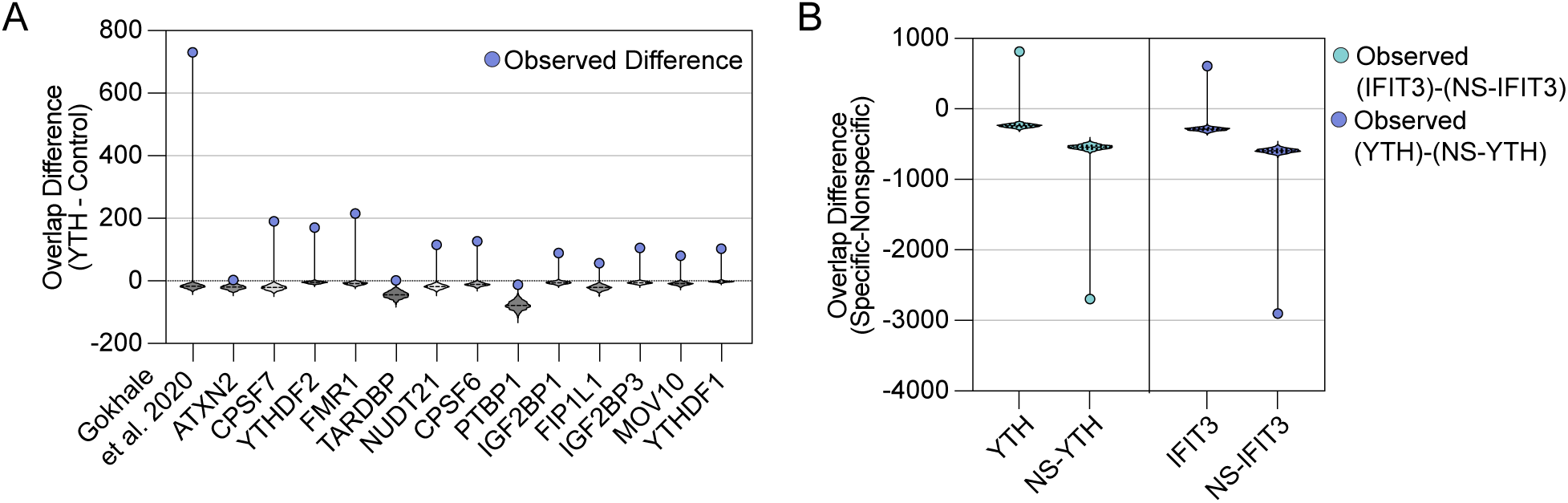
Validation of ADAR-YTH editing specificity and IFIT3-ADAR/ADAR-YTH overlap analyses, related to Figure 2. (**A**) Permutation test distributions (1,000 shuffles) for overlap of ADAR-YTH–specific or non-specific edit sites with meRIP-seq peaks and indicated POSTAR3 CLIPdb datasets. The observed overlap difference (YTH − Control) is indicated for each comparison. (**B**) Permutation test distributions showing the difference in overlap between specific and non-specific edit sites for ADAR-YTH (YTH − NS-YTH) and IFIT3-ADAR (IFIT3 − NS-IFIT3). Observed differences are indicated by vertical lines.

We found that IFIT3-ADAR edits significantly overlap with ADAR-YTH edits (100 bp window, 1,000 shuffles, FDR<0.005), but not with non-specific ADAR-YTH edits that did not pass our filtering criteria (**Figure 2B & D, Supplementary Figure 2B**). Reciprocal analysis showed that ADAR-YTH directed-edits also overlap with IFIT3-ADAR edits, whereas randomly shuffled IFIT3-ADAR control sites showed no enrichment (**Figure 2D, Supplementary Figure 2B**). Notably, we identified four ADAR-YTH edits within the HCV genome that overlap with IFIT3-ADAR edits (**Figure 2E**), consistent with m^6^A-dependent IFIT3 binding to HCV RNA (**Figure 2A**). Treatment of cells with the METTL3 inhibitor STM2457 reduced both ADAR-YTH and IFIT3-ADAR editing at HCV sites 2098 and 3131 over 50% relative to the control, whereas site 6586 remained unaffected (**Figure 2F**). These findings establish m^6^A as a determinant of IFIT3 association with both cellular and HCV RNA.

### IFIT3•RNA interaction is enhanced by m^6^A and requires cellular co-factors and *cis*-acting viral sequences

Having established that IFIT3 preferentially associates with m^6^A-modified transcripts, we next defined biochemical requirements for this interaction. In streptavidin pulldown assays using biotinylated 38-nucleotide probes containing a single m^6^A or an unmodified adenosine (**Figure 3A**), IFIT3-V5 from IFN-β-stimulated 293T lysates co-purified preferentially with the m^6^A-modified probe (**Figure 3B**). Purified recombinant IFIT3 did not bind any of these probes (**Figure 3C, Supplementary Figure 3A**). Control proteins confirmed the integrity of the assay: recombinant YTHDC1 preferentially bound m^6^A-modified probes (**Figure 3D**), and recombinant IFIT1 bound the 5’-triphosphate-containing probes regardless of m^6^A status (**Supplementary Figure 3B**). The discrepancy between lysate-derived and purified IFIT3 therefore indicates that IFIT3•RNA association requires additional cellular co-factors.

**Figure 3.**
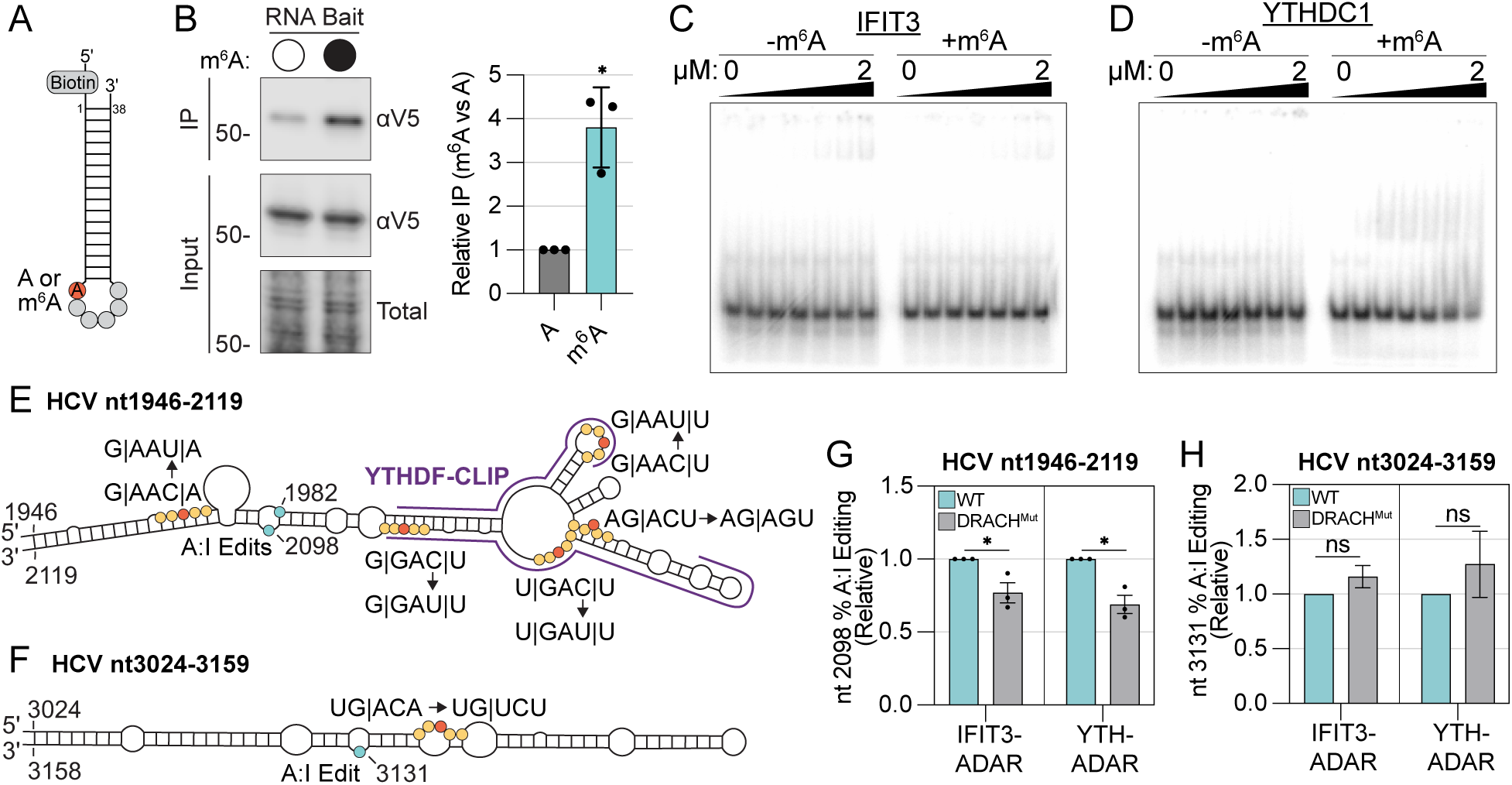
IFIT3•RNA interaction is enhanced by m⁶A and requires cellular co-factors and *cis*-acting viral sequences. (**A**) Schematic of biotinylated RNA probe +/- m⁶A used for cell-free RNA binding assays. (**B**) Representative immunoblot and corresponding quantitation across replicates of IFIT3-V5 from IFNβ-treated (100 U/ml, 24 h) 293T cytoplasmic extracts bound to synthetic biotinylated RNA probes after streptavidin-bead purification and washing. Lanes represent independent RNA affinity pulldown assays where +/- m⁶A probes were incubated with extracts. (**C-D**) Electrophoretic mobility shift assay of IFIT3 and YTHDC1 with RNA probe (as in A with no 5’ biotin) ± m^6^A modification. For all gels, protein concentrations in each well are: 0, 0.065, 0.130, 0.260, 0.521, 1.04 and 2.08 μM (left to right). (**E**) Diagram of DRACH motif mutations introduced into HCV genomic RNA in proximity to the 2098 IFIT3-ADAR and YTH-ADAR editing sites (indicated as cyan dot). Previous SHAPE-seq^47^ was used to depict secondary structure. Published YTHDF PAR-CLIP peaks^43^ are indicated in purple. Yellow and red dots represent DRACH motifs with red dots indicating putative m^6^A nucleotides. Nucleotide mutations are indicated at respective DRACH motifs with codons separated by “|”. (**F**) Diagram of DRACH motif mutations introduced into HCV genomic RNA in proximity to the 3131 IFIT3-ADAR and YTH-ADAR editing sites (indicated as cyan dot). Annotations are as explained in (E). (**G**) RT-PCR Sanger sequencing quantification of IFIT3-ADAR and YTH-ADAR A:I editing at nt 2098 on WT or mutant HCV RNA 72 h post electroporation. Values are normalized to WT editing. (**H**) RT-PCR Sanger sequencing quantification of IFIT3-ADAR and YTH-ADAR A:I editing at nt 3131 on WT or mutant HCV RNA 72 h post electroporation. Values are normalized to WT editing. See also Figure S3.

**Supplementary Figure 3.**
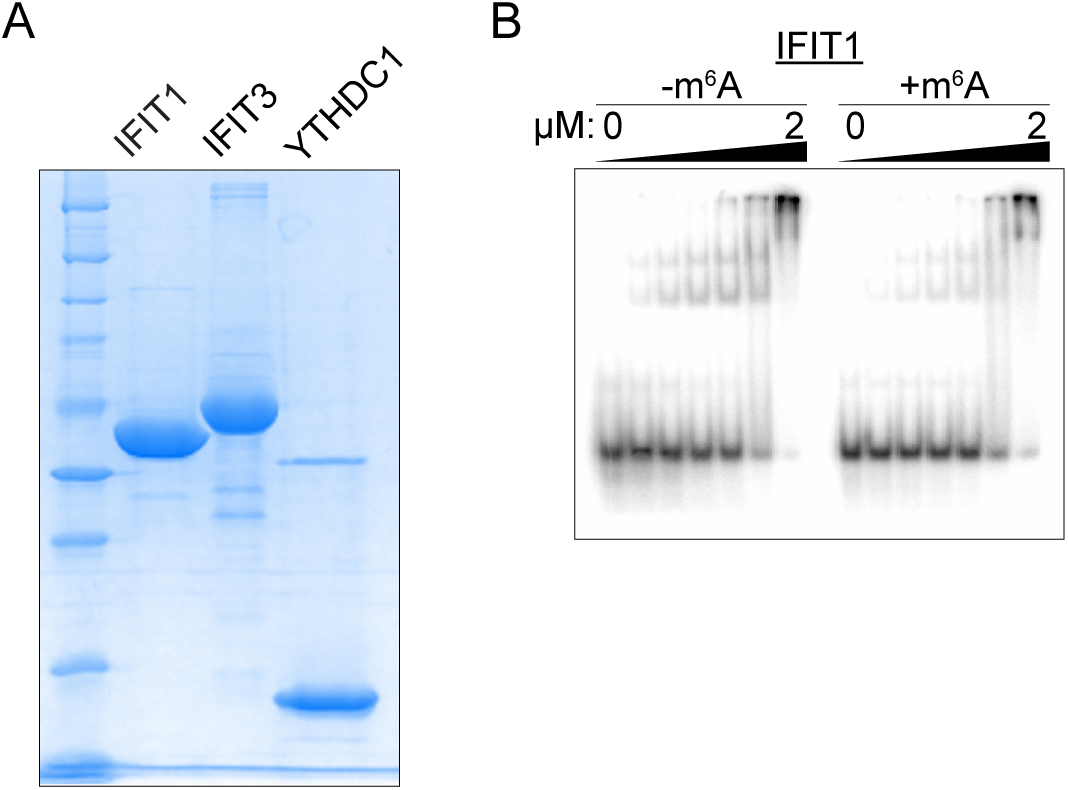
IFIT3 protein purification, related to Figure 3. (**A**) SDS–PAGE analysis of the indicated purified recombinant proteins, visualized by Coomassie Blue staining. (**B**) Electrophoretic mobility shift assay of IFIT1 with RNA probe (as in A with no 5’ biotin) ± m^6^A modification. For all gels, protein concentrations in each well are: 0, 0.065, 0.130, 0.260, 0.521, 1.04 and 2.08 μM (left to right).

We next asked whether m^6^A on viral RNA similarly promotes IFIT3 recognition in cells. To test this, we generated full-length HCV RNA molecules carrying synonymous mutations (where possible) in DRACH motifs near IFIT3-ADAR & YTH-ADAR edits (**Figure 3E-F)**^43^. Mutational strategy and target sites were selected based on published SHAPE-seq and YTHDF PAR-CLIP-seq data indicating that DRACH motifs surrounding the IFIT3–ADAR edit site at 2098 are amenable to mutation and likely m^6^A-modified^43,47^. As a specificity control, the DRACH motif near edit site 3131, which had no orthogonal evidence of m^6^A modification, was also mutated, as it is possible that distant m^6^A sites drive editing. We transfected purified wild-type or DRACH-mutant HCV RNA into cells expressing IFIT3-ADAR and YTH-ADAR and measured editing proximal to the mutated motifs at 48 hours post-transfection. Mutation of all five DRACH motifs within nucleotides 1946-2110 reduced both IFIT3-ADAR and YTH-ADAR editing at site 2098 (**Figure 3G**), whereas mutation of the DRACH motif near site 3131 had no effect (**Figure 3H**). These data demonstrate that m^6^A-containing sequences within the HCV genome directly contribute to IFIT recognition of viral RNA.

### IFIT3•RNA association is mediated by a predicted helical hairpin independent of IFIT1/2 interaction

Because our data suggest that IFIT3•RNA interaction may require cellular cofactors, we next tested which IFIT3 domains mediate RNA binding, focusing on whether interactions with IFIT1 and IFIT2 are involved. Using RNA affinity pulldown assays with m^6^A-containing probes, we tested a panel of deletion mutants expressed in IFNβ-treated 293T IFIT3 knockout cells (**Figure 4A-C**). We identified two regions critical for RNA association: TPRs 1-2 and the uncharacterized region (UCR; residues 274-415) between TPRs 6-7 (**Figure 4B-C**). TPRs 1-2 align with a recently characterized domain swapping interaction, termed the “swap domain,” between murine IFIT2 and IFIT3^12^ while the UCR overlaps with residues required for IFIT3-dependent enhancement of influenza A virus translation^11^, as modeled by AlphaFold3 (UniProt: O14879; **Figure 4D**)^49^. To determine where these RNA-binding domains also mediate protein-protein interactions, we performed co-immunoprecipitation assays in IFIT1-IFIT3 or IFIT1-IFIT2 double-knockout 293T cells. All RNA-binding deletion mutants retained IFIT1 interaction, including the UCR deletion; only removal of the known IFIT1-binding interface (residues 403-490)^16^, abolished it (**Figure 4E**). In contrast, deletions of TPRs 1–2 or 3–4, but not the UCR, disrupted IFIT2-IFIT3 association (**Figure 4F**). Thus, TPRs 1–2 and the UCR are both required for RNA binding, but only the UCR supports RNA binding without affecting IFIT3 interactions with IFIT1 or IFIT2.

**Figure 4.**
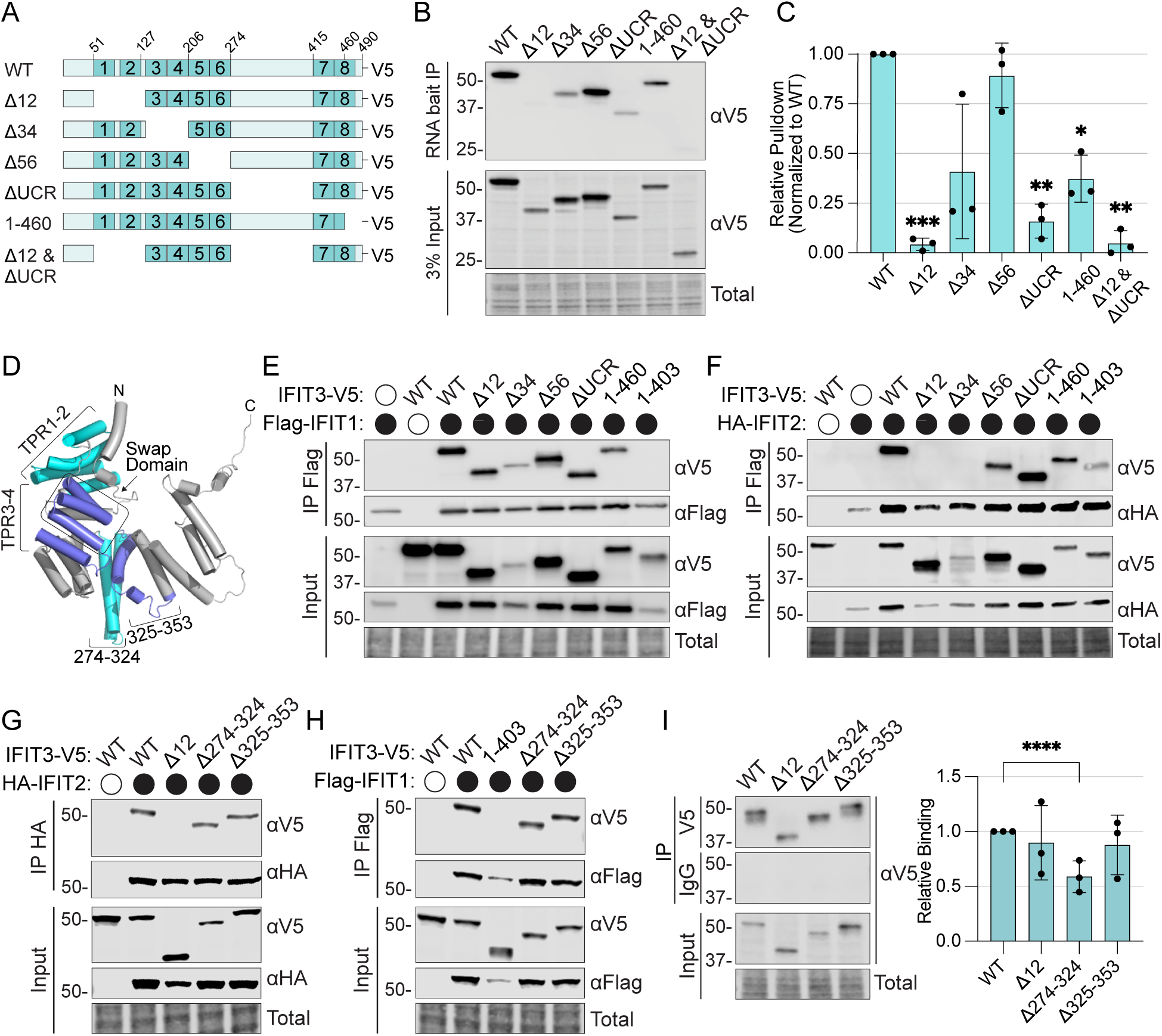
IFIT3•RNA association is mediated by a predicted helical hairpin independent of IFIT1/2 interaction. (**A**) Diagram of human IFIT3 domain deletions used for assays B-I. (**B-C**) Representative immunoblot and corresponding quantitation across replicates of transfected IFIT3-V5 WT or mutant (see 4A) from IFNβ-treated (100 U/ml, 24 h) IFIT3 knockout 293T cytoplasmic extracts bound to synthetic biotinylated RNA probes after streptavidin-bead purification and washing. Lanes represent independent RNA affinity pulldown assays where m⁶A probes (see 3A) were incubated with extracts. (**D**) Predicted 3d structure of human IFIT3 (UniProt: O14879, Alphafold3 server^48^) with TPRs, putative IFIT2 interaction “swap domain”^12^, and target regions for deletion annotated. (**E**) Immunoblotting of IFIT3-V5 WT or mutant (see 4A) co-precipitation with Flag-IFIT1 co-expressed in IFNβ-treated (100 U/ml, 24 h) IFIT1-IFIT3 double-knockout 293T cells. (**F**) Immunoblotting of IFIT3-V5 WT or mutant (see 4A) co-precipitation with HA-IFIT2 co-expressed in IFNβ-treated (100 U/ml, 24 h) IFIT1-IFIT2 double-knockout 293T cells. (**G**) Immunoblotting of IFIT3-V5 WT or mutant (see 4D) co-precipitation with Flag-IFIT1 co-expressed in IFNβ-treated (100 U/ml, 24 h) IFIT1-IFIT3 double-knockout 293T cells. (**H**) Immunoblotting of IFIT3-V5 WT or mutant (see 4D) co-precipitation with HA-IFIT2 co-expressed in IFNβ-treated (100 U/ml, 24 h) IFIT1-IFIT2 double-knockout 293T cells. (**I**) Representative Immunoblotting of expression and immunoprecipitation of stably expressed WT or mutant IFIT3-V5 in HCV-infected (MOI 1, 24 hpi) IFIT1-IFIT3 double-knockout Huh7 cells for native RIP-qPCR (left). Native RIP-qPCR of co-purified HCV RNA from immunoprecipitated fractions shown in. Values represent enrichment of V5-purified RNA in comparison to IgG followed by normalization to WT IFIT3 (right).

To define the contribution of the UCR in more detail, we generated smaller deletions targeting residues 274–324 or 325–374. These segments form a helical hairpin in mouse IFIT3 and are predicted to adopt the same fold in human IFIT3 (**Figure 4D**)^12,48^. Consistent with the larger UCR deletion, neither subdomain deletion disrupted IFIT1 and IFIT2 interaction (**Figure 4G-H**). We then tested for whether these mutants could associate with HCV RNA in cells. Because RNA binding could be abrogated without disrupting IFIT1 interaction (**Figure 4E**), we stably expressed these constructs in Huh7-IFIT1/IFIT3 double-knockout cells to avoid any indirect effects of IFIT1 on IFIT3 RNA binding. RIP-qPCR revealed that deletion of residues 274–324, which does not affect IFIT1 or IFIT2 interaction, reduced IFIT3 pulldown of HCV RNA, whereas the ΔTPR1-2 mutant, which disrupts IFIT2 interaction, did not (**Figure 4G-I**). Because ΔTPR1-2 retained HCV RNA binding in cells despite loss of short probe binding in cell-free assays (**Figure 4B–C**), these data indicate the UCR to be critical for RNA binding while TPRs 1-2 may influence binding indirectly through protein-protein interaction and/or specific RNA contexts. These findings identify a conserved helical hairpin within the UCR as essential for IFIT3 recognition of HCV RNA in cells, independent of IFIT1 and IFIT2 interactions.

### IFIT3 recognition of HCV RNA limits viral RNA release

Having defined the domain requirements for IFIT3–HCV RNA interaction, we next tested whether these domains also contribute to IFIT3-mediated restriction of HCV infection. We stably expressed V5-tagged wild-type IFIT3, ΔTPR1–2 (disrupts IFIT2 interaction), or Δ274–324 (disrupts HCV RNA binding) in Huh7 cells and infected them with HCV. At 48 hpi, HCV protein abundance (**Figure 5A**) and intracellular HCV RNA levels (**Figure 5B**) were unchanged across cell lines. In contrast, HCV RNA measured in clarified culture supernatants was reduced relative to the parental control (set to 100%) to 57% in wild-type IFIT3 cells and 82% in ΔTPR1–2 cells (**Figure 5C**), whereas Δ274–324 showed no significant change. These results indicate that IFIT3 reduces released/supernatant-associated HCV RNA and that this activity requires an IFIT3 RNA-binding interface.

**Figure 5.**
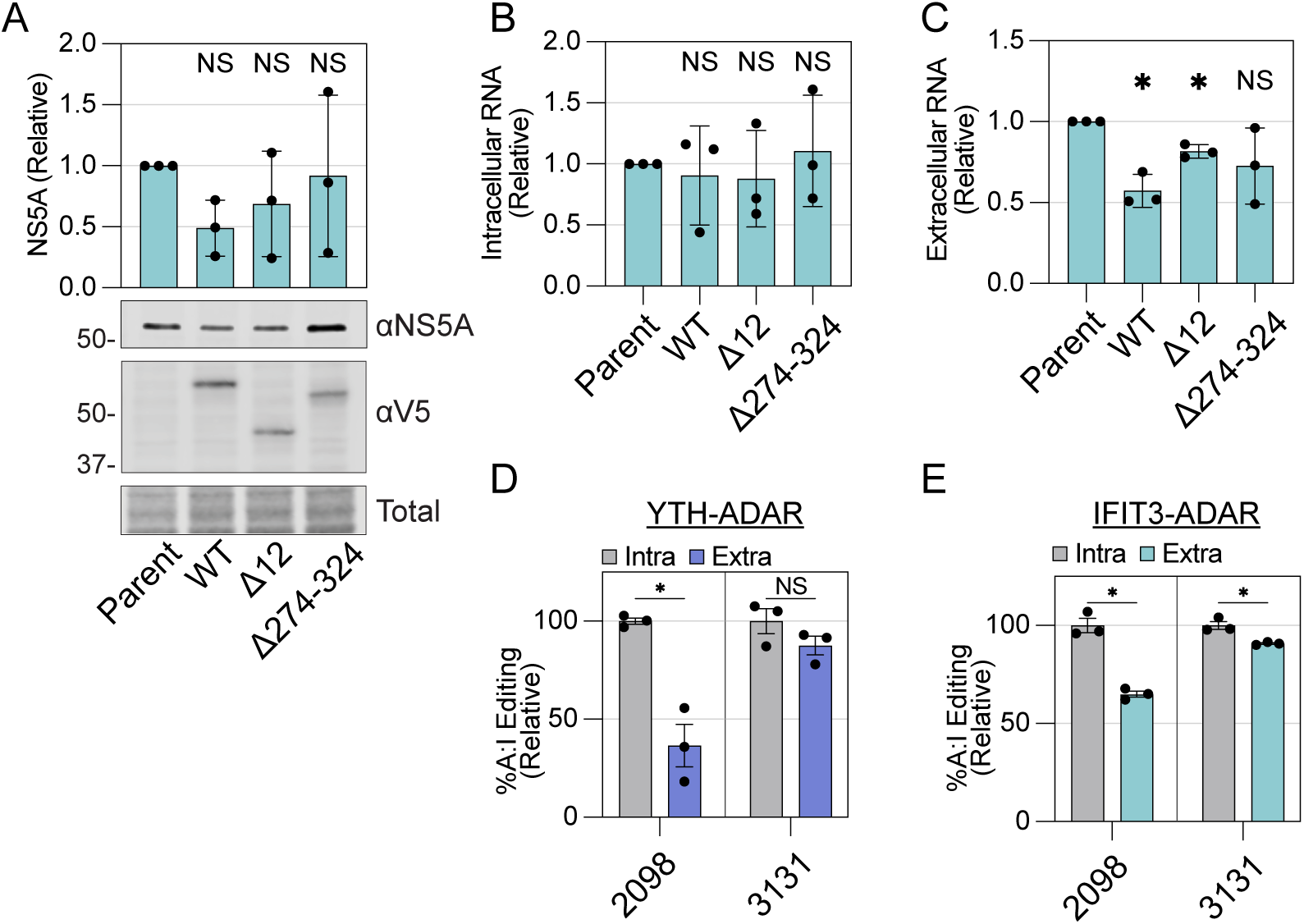
IFIT3 recognition of HCV RNA limits viral RNA release. (**A**) Quantification of HCV NS5A immunoblotting in stably-expressed WT or mutant IFIT3 in HCV-infected (MOI 1) Huh7 cells at 48 hpi and representative blot of the HCV NS5A protein and IFIT3-V5. (**B**) Relative RT-qPCR quantification of intracellular HCV RNA copies from (A) normalized to Parent cell lines at 48 hpi. (**C**) Relative RT-qPCR quantification of extracellular HCV RNA copies from (A) normalized to Parent cell lines at 48 hpi. (**D**) RT-PCR Sanger sequencing quantification of YTH-ADAR A:I editing on intracellular and extracelluar HCV RNA at indicated sites. Values are normalized to intracellular editing. (**E**) RT-PCR Sanger sequencing quantification of IFIT3-ADAR A:I editing on intracellular and extracelluar HCV RNA at indicated sites. Values are normalized to intracellular editing.

m^6^A has been proposed to restrict late stages of the HCV life cycle, including reduced viral output upon overexpression of YTHDF m^6^A readers and lower m^6^A levels on extracellular HCV RNA compared with intracellular viral RNA^43^. As our data establishes an m^6^A preference for IFIT3 RNA binding, we hypothesized that IFIT3 restricts infection similary. As a benchmark for this model, we compared YTH-ADAR editing on intracellular HCV RNA versus the viral RNA in the supernatant and observed reduced editing at site 2098 in the supernatant fraction (**Figure 5D**). We then asked whether IFIT3 shows a similar bias. IFIT3-ADAR editing at site 2098 was likewise reduced in supernatant HCV RNA (**Figure 5D**). IFIT3-ADAR editing at site 3131 was also reduced in supernatant RNA, whereas YTH-ADAR showed no corresponding decrease at this site (**Figure 5D-E**). Together, these data suggest that IFIT3 preferentially associates with an intracellular pool of HCV RNA and limits viral output in a manner resembling YTHDF proteins.

## DISCUSSION

Structural and biochemical studies have defined cap0 recognition by IFIT1, AU-rich element binding by IFIT2, and diverse RNA binding by IFIT5, but the RNA-binding properties of IFIT3 have remained poorly understood (reviewed in Mears and Sweeney^7^)^14,16,19,21,22,49,50^. Recent work has begun to address this gap: Sullivan et al. showed that IFIT3 directly binds influenza A virus RNA and enhances viral mRNA translation, while Glasner et al. demonstrated that murine IFIT2 and IFIT3 form a heterodimer that restricts negative-strand viruses by targeting mRNAs with short 5′ UTRs^11,12^. Our findings extend this emerging picture by providing a transcriptome-wide map of IFIT3 RNA targets on host and viral RNA during HCV infection. Systematic comparison of these targets to published RBP binding profiles revealed that IFIT3 shares binding preferences with a variety of m^6^A reader proteins, an overlap that led to the identification of m^6^A as a previously unrecognized determinant of IFIT protein-RNA interaction.

The identification of m^6^A as an IFIT3-binding determinant is supported by four independent lines of evidence: transcriptome-wide overlap of IFIT3-ADAR editing sites with the previously established m^6^A-sensor YTH-ADAR construct^29^, loss of IFIT3•RNA association upon METTL3 inhibition, preferential co-purification with m^6^A-modified probes in cell-free assays, and reduced IFIT3-ADAR editing at HCV sites when *cis*-acting DRACH motifs were mutated on the viral genome itself. These data establish that IFIT3•RNA association depends on m^6^A but do not exclude an indirect model in which canonical m^6^A readers recruit IFIT3 to methylated sites. However, the requirement for the UCR helical hairpin (274-324), a domain dispensable for all tested protein–protein interactions, argues that IFIT3 likely contributes an RNA-contact surface. Although both IFIT2 and IFIT3 bind preferentially in 3′ UTRs, IFIT3 targets did not overlap with published IFIT2 RNA-binding datasets, reinforcing that m^6^A status distinguishes IFIT3 specificity from that of its closest family member. We found that IFIT3 and YTHDF proteins share binding sites on HCV RNA, raising the question of whether IFN-induced IFIT3 competes with constitutively expressed YTHDF readers for m^6^A-marked viral RNA. Such competition could shift the functional consequences of m^6^A on HCV RNA from YTHDF-mediated regulation of packaging and translation to IFIT3-mediated restriction, effectively repurposing a host RNA modification as an antiviral signal under interferon conditions.

Domain mapping experiments reveal that IFIT3•RNA interaction is structurally separable from its cofactor roles. TPRs 1–2 and residues 274–324 both contribute to RNA association on short probes, but only the helical hairpin is required for HCV RNA binding in cells, a distinction consistent with the UCR serving as the primary RNA-recognition element while TPR1-2 provides auxiliary function either through protein-protein interaction or potentially contacting RNA. Importantly, deletion of the helical hairpin does not disrupt IFIT1 or IFIT2 interaction indicating that IFIT3 contains a dedicated RNA-binding surface distinct from the C-terminal IFIT1-interaction domain and the N-terminal IFIT2-swapped interface^12,16^. Sullivan et al. similarly mapped an independent RNA-binding surface on IFIT3, mediated by several residues within amino acids 249-286, that engages RNA without disrupting IFIT heterocomplexes^11^. Together, these findings from two independent studies converge on a model in which IFIT3 harbours autonomous RNA-binding capacity on surfaces separate from its protein-protein interaction domains.

How m^6^A enhances IFIT3•RNA association remains an open question. Although IFIT3 preferentially co-purified with an m^6^A-modified probe in cell extracts, purified recombinant IFIT3 did not bind this same probe *in vitro*, in contrast to purified IFIT1, which bound 5′-triphosphate probes as expected. This discrepancy may reflect a requirement for post-translational modifications, chaperone-assisted folding, or an accessory factor present in cellular extracts. Although Sullivan et al. detected binding of recombinant IFIT3 to longer (∼200 nt) RNA probes^11^, raising the possibility that probe length and RNA sequence are key variables to IFIT3 RNA association. Given that TPR1-2 deletion, which disrupts IFIT2 interaction, reduced binding to short probes in our assays, it is likely that IFIT3 RNA requires cofactors for certain RNA binding contexts. This was observed by Glasner et al. who showed that eCLIP-seq of murine IFIT3, a measure of direct RNA protein interaction, required co-expression of IFIT2 for detectable binding^12^. Nonetheless, resolving whether IFIT3 is a direct m^6^A reader or an m^6^A-dependent RNA-binding protein that requires additional cellular context will be an important area for future investigation.

Functionally, IFIT3 restriction of HCV required its RNA-binding capacity and resembled previously established models for m^6^A-dependent viral restriction. Notably, IFIT3 restricted extracellular but not intracellular HCV RNA, raising the possibility that IFIT3•RNA binding affects viral RNA packaging and virion production rather than earlier viral lifecycle stages. Consistent with this model, IFIT3-ADAR editing at HCV site 2098 and 3131 was reduced on extracellular relative to intracellular viral RNA, indicating IFIT3-targeted RNA is specifically prevented from exiting the cell. This is consistent with previous m^6^A-dependent antiviral models where m^6^A-modified HCV RNA was not efficiently packaged into virions^43^, a result replicated in this study by showing extracellular HCV RNA had less YTH-ADAR editing that intracellular. Whether IFIT3 directly blocks HCV RNA interaction with the Core packaging protein, as shown with m^6^A^43^, or acts through an as-yet-unidentified mechanism remains to be determined.

Beyond viral RNA, IFIT3 bound a broad set of host transcripts enriched for membrane-associated compartments, including the ER and Golgi apparatus. Because HCV replication and assembly occur on ER-derived membranes^51^, the overlap between IFIT3-bound host mRNAs and the sites of viral biogenesis raises the possibility that IFIT3 modulates the local environment at replication organelles. Whether IFIT3 binding to these host transcripts alters their stability or translation, and whether this contributes to antiviral defense or reflects a broader role in regulating membrane-associated gene regulation during the IFN response, remains to be determined.

These findings establish that an ISG-encoded effector can indeed exploit m^6^A as a recognition determinant for antiviral restriction. Rather than acting exclusively as a cofactor that tunes IFIT1 or IFIT2 RNA-binding activity, IFIT3 functions as an autonomous factor whose RNA preferences are shaped by the m^6^A landscape of the cell. This is particularly relevant given that m^6^A marks the RNA genome of many viruses beyond HCV^52^. More broadly, our data add to an emerging view that the IFN response reshapes not only the repertoire of expressed RBPs but also the RNA-binding activity of those already present in the cell^53–55^. The observation that recombinant IFIT3 does not bind short m^6^A-containing probes, whether because it requires cellular cofactors, IFN-induced post-translational modifications, or simply longer RNA substrates, suggests that the intracellular environment plays a key role in licensing IFIT3•RNA engagement. The consequences of m^6^A for infection may therefore be context-dependent: the same RNA modification that regulates viral particle production could also flag viral RNA for IFIT3-mediated restriction when a functional IFN response is active. This is notable given that m^6^A on viral RNA has primarily been described as preventing recognition by RIG-I and MDA5, thereby enabling immune evasion^56,57^. Our findings suggest that while m^6^A may shield viral RNA from these cytoplasmic sensors, it simultaneously renders that RNA visible to a distinct class of IFN-induced effector. Whether this mechanism extends to other m^6^A-modified viruses, and how IFIT3’s autonomous and complex-dependent activities are balanced across different viral contexts, are important questions for future investigation.

## DATA AND CODE AVAILABILITY

The RNA-seq data generated in this study will be deposited in NCBI Gene Expression Omnibus (GEO) prior to journal publication; the accession number will be provided once available. Previously published data used in this study was obtained by CLIPdb bulk download^33^ or from supplementary bed files for meRIP-seq^46^ and IFIT2 CLIP-seq^14,32^. All original code and additional information required to reanalyze the data reported in this paper is available from the lead contact upon request.

## ACKNOWLEDGMENTS

We thank current and past members of the Horner Lab for valuable feedback and discussion. Research on m^6^A in the Horner lab has been supported by Burroughs Wellcome Fund and National Institutes of Health (NIH) grant R01AI125416. MGT has received support from NIH T32CA009111 and the American Cancer Society. Y.N was supported by the National Institutes of Health (R01CA258589 to Y.N.), the Welch Foundation (I-2115--20220331 to Y.N.), and the Cancer Prevention and Research Institute of Texas (CPRIT training grant RP210041 to N.S.).

## DECLARATION OF INTERESTS

The authors declare that they have no conflicts of interest with the contents of this article. The content is solely the responsibility of the authors and does not necessarily represent the official views of the National Institutes of Health

## METHOD DETAILS

### Cell Culture

All cell lines were grown in Dulbecco’s modified Eagle’s medium (DMEM; Mediatech) supplemented with 10% fetal bovine serum (FBS; HyClone), 25 mM HEPES (Thermo Fisher), and 1× non-essential amino acids (Thermo Fisher), referred to as complete DMEM (cDMEM). Huh7 and Huh-7.5 cells were verified using the Promega GenePrint STR kit (DNA Analysis Facility, Duke University) and confirmed mycoplasma free by the LookOut Mycoplasma PCR Detection Kit (Sigma-Aldrich).

### Plasmids

The following previously published plasmids were used in this study: pMD2.G (RRID: Addgene_12259), psPAX2 (RRID: Addgene_12260), pLentiCRISPRv2-IFIT3^14^, and psJFH-1-M9^58^. Other plasmids were generated using standard cloning techniques or mutational PCR followed by multi-segment ligation as recommended in the InFusion HD Cloning Kit (Takara). pLVX-derived plasmids were generated by insertion of PCR amplicons into pLVX-EF1α-eGFP-2xStrep-IRES-Puro (RRID: Addgene_141395) at restriction sites downstream of the EF1α promoter and upstream of the IRES-Puro element. Hygromycin resistance was added to pLVX plasmids containing WT or mutant IFIT3 by removing the Puromycin resistance gene with restriction enzyme digestion and replacing with a Hygromycin resistance gene. TLCV2-derived plasmids were generated by insertion of PCR amplicons into TLCV2-APOBEC1-YTH (RRID: Addgene_178949) at restriction sites downstream of the Tet-On promoter and upstream of the EF1α-Puro element. TRIBE-seq ADAR constructs were derived by insertion of the ADAR catalytic domain from pCMV-ADARcd-YTHD422N (RRID: Addgene_194402) into the indicated vectors. IFIT-derived plasmids were generated from pEF1-IFIT1-V5, pEF1-IFIT2-V5, and pEF1-IFIT3-V5^14^. DRACH Mutant HCV plasmids were produced by insertion of a gBlock containing relevant mutations between AgeI and SpeI restriction enzyme sites (psJFH-1-M9-sl1946mut) or by mutational PCR followed by InFusion cloning (psJFH-1-M9-sl3020mut). Human YTHDC1 (amino acids 345 to 509) was cloned into pET-21(+) (Novagen) with an N-terminal hexahistidine tag and a TEV cleavage site. The full-length human IFIT1 gene was cloned into pSMT3 (gift of Chris Lima) with an N-terminal hexahistidine tag, thrombin cleavage site, and SUMO tag. The full-length human IFIT3 was cloned into pET15b with an N-terminal hexahistidine tag and a TEV cleavage site. All plasmids were validated by sequencing.

### Viruses

HCV infectious stocks (cell culture-adapted strain of genotype 2A JFH1 HCV) were generated in Huh-7.5 cells and quantified by focus-forming assay (FFA), as described^58^. Viral genomic sequences were validated by long-read sequencing (Plasmidsaurus) of full-length complementary DNA (cDNA) amplicons. Unless otherwise noted, cells were inoculated with HCV (indicated by multiplicity of infection (MOI)) diluted in serum-free DMEM for 3 h. After inoculation, virus was removed and replaced with cDMEM.

### Generation of Cell Lines

Lentiviral stocks were generated in 293T cells transfected with lentiviral vectors and pMD2.G and psPAX2 using TransIT-LT1 transfection reagent (Mirusbio). Target cells were transduced by these lentiviruses and stable cell lines selected with 2 µg/ml Puromycin (Sigma-Aldrich) to generate Huh7 IFIT3-ADAR, Huh7 YTH-ADAR, Huh7 ADAR-Ctl, Huh7 IFIT3. To generate Huh7 IFIT1/IFIT3 double-knockout + WT IFIT3, Huh7 IFIT1/IFIT3 double-knockout + Δ12 IFIT3, and Huh7 IFIT1/IFIT3 double-knockout + ΔUCR IFIT3, following lentiviral transduction, cells were selected with 250 µg/ml Hygromycin B (Thermo Fisher) to generate pools that were used immediately. To generate Huh7 IFIT3 knockout cells, lentiviral stocks were generated using pLentiCRISPRv2-IFIT3, as above, and following transduction, stable cell lines were selected with 2 µg/ml Puromycin (Sigma-Aldrich). Clones were screened by immunoblot and Sanger sequencing, analyzed with the EditCo CRISPR ICE tool. For electroporation-based knockouts of IFIT1 or IFIT2, multi-guide sgRNA and recombinant Cas9 (EditCo) were incubated to form RNP complexes and electroporated into parental Huh7 or 293T WT or IFIT3 knockout cells using a Neon Transfection System (Life Technologies; 1,200 V, 50 ms, 1 pulse) according to the Synthego Gene Knockout Kit v2 protocol. Knockout efficiency was assessed by immunoblot and Sanger sequencing (EditCo CRISPR ICE) of genomic DNA-derived amplicons.

### *In vitro* transcription and electroporation of HCV RNA

HCV RNA was *in vitro* transcribed from XbaI linearized WT or DRACH mutant psJFH-1-M9 plasmids using the MEGAscript T7 transcription kit (Thermo Fisher), according to manufacturer protocol. Electroporation into Huh7 WT or mutant cell lines was performed using previously described methods^59^. 6×10^6^ cells resuspended in 400 µl cold cytomix buffer (120 mM KCl; 0.15 mM CaCl_2_; 10 mM K_2_HPO_4_/KH_2_PO_4_, pH 7.6; 25 mM HEPES, pH 7.6; 2 mM EGTA, pH 7.6; 5 mM MgCl_2_, pH adjusted with KOH; 2 mM ATP and 5 mM glutathione) were electroporated with 5 µg of *in vitro* transcribed HCV RNA in a 4 mm cuvette using a BioRad GenePulser XCell with CE module (BioRad; 270 V, 950 µF). Cells were immediately added to 13 ml cDMEM and incubated for 10 minutes at RT. 1 ml of cell suspension was plated in 6-well plates for followed by PBS washing and fresh media 6 h post-plating.

### Cell Treatments

For METTL3 inhibitor experiments, cells were treated with 20 µM STM2457 (MedChemExpress) freshly reconstituted in dimethyl sulfoxide (DMSO), or an equivalent volume of DMSO. After 72 h, cells were replated in STM2457- or DMSO-containing media for subsequent experiments. For IFN treatments, IFNβ stock (100 U/µl in PBS; PBL) was diluted to the indicated final concentration in cDMEM before being added to cells.

### HyperTRIBE-seq and Analysis

Huh7-IFIT3-ADAR-V5, Huh7-ADAR-YTH-HA, or Huh7-ADAR-V5 pools were infected with HCV (MOI 1). Post-inoculation, virus was replaced with cDMEM supplemented with 0.5 U/ml IFN-β1 (PBL). At 72 h post-infection, RNA was purified from two-thirds of cells using TRIzol (Thermo Fisher) according to the manufacturer’s protocol; the remainder was used for immunoblot verification of fusion protein expression. Library preparation and sequencing were performed at Azenta Life Sciences (standard Illumina RNA-seq with ribosomal RNA (rRNA) depletion, 2 × 150 bp, ∼350M paired-end (PE) reads across 9 samples).

Raw FASTQ files were processed on the Duke Compute Cluster using nf-core/rnaseq v3.14.0^60^ within the nf-core framework^61^, with reproducible software environments from Bioconda^62^ and BioContainers^63^, executed with Nextflow v23.10.1^60^. Reads were first mapped to the human GRCh38 reference genome; unmapped reads were subsequently aligned to the HCV genome (GenBank: AB047639.1) using the same workflow. RNA editing rates were quantified using the Bullseye pipeline^37,38^ with modifications (https://github.com/mflamand/Bullseye/). Count matrices were generated from aligned BAM files using parsebam.pl for each sample. A-to-G transition frequencies were calculated using Find_edit_site.pl, comparing each test sample against 3 replicates of the reference dataset (e.g., IFIT3 vs. ADAR). Only sites with >1% A-to-G editing and ≥10 total counts in all replicates of both conditions were considered. Genomic coordinates of edit sites were overlapped using bedtools intersect, and editing frequencies in intersecting, unique, and combined sets were plotted for qualitative assessment. Significant differences in editing between conditions were tested using a quasibinomial generalized linear model in followed by Benjamini–Hochberg false discovery rate (FDR) adjustment using R. Results were filtered for the intersecting subset at adjusted *p* < 0.05 before plotting log₂ fold change (LFC) and editing rates. A base editing cutoff of 8% was chosen based on the relationship between LFC values and editing rates, applied to all samples; the LFC cutoff was applied when editing was quantified in both conditions. Cutoffs were determined from host data and applied to viral data post hoc. All original analyses code is available upon request.

### Functional annotation/GO analyses

The DAVID functional annotation tool was used for GO analyses of genes of interest^35,64^. For GO analyses of HyperTRIBE-seq data, a gene list derived from the union of all genes edited by IFIT3-ADAR, ADAR-YTH, and ADAR-V5 served as the background data set. For GO analyses of RBPs overlapping with HyperTRIBE-edited genes, a list of all RBPs tested in CLIPdb was used as a background. Default settings were used for statistical testing and p-value adjustment. Functional annotation charts were exported and plotted with GraphPad Prism.

### Permutation Analyses of HyperTRIBE-seq

To test whether IFIT3-ADAR or YTH-ADAR editing sites overlap a comparison dataset more than expected from non-specific ADAR editing, we performed a windowed overlap analysis with an empirical permutation test (100 nt window) using a combination of Python scripts and the bedtools package^65^. IFIT3-ADAR, YTH-ADAR, and matched ADAR-only control BED files were converted to strand-specific pseudo-peaks by clustering sites on the same strand (bedtools cluster, max inter-site distance 100 nt), merging clustered intervals (bedtools groupby), and expanding by 100 nt (bedtools slop).

Overlaps with the comparison dataset were quantified using bedtools intersect (-u -s) and the test statistic was defined as Δ = overlaps_test_ – overlaps_control_. Significance was assessed by independently shuffling the test and control pseudo-peaks within a whitelist of expressed gene intervals (preserving interval lengths and chromosome boundaries) and recomputing Δ to generate a null distribution (1,000 permutations). Empirical p-values were computed as a right-tailed permutation *p*-values with +1 correction, p = (1 + #(Δ*_perm_* ≥ Δ*_obs_*))/(1 + *N_perm*) and followed by Benjamini–Hochberg FDR adjustment.

### Quantitative RT-PCR

Purified RNA was reverse transcribed using the iScript cDNA Synthesis Kit (Bio-Rad). cDNA was diluted 1:5, and quantitative PCR (qPCR) was performed in technical triplicate using Power SYBR Green PCR Master Mix (Thermo Fisher) on a QuantStudio 6 Flex or QuantStudio Pro system (Thermo Fisher). Raw C_t_ values were exported and analyzed by the ΔΔCt method.

### Viral RNA isolation and copy number quantification

Intracellular and extracellular RNA were isolated using TRIzol and TRIzol LS, respectively (Thermo Fisher). After liquid-liquid extraction, the aqueous fraction was combined with 4 µg yeast tRNA (Invitrogen) and 1.5 volumes 100% EtOH, then DNase treated and eluted in 20 µL using Zymo RNA Clean & Concentrator-5 columns (Zymo Research). Viral copy numbers were determined by standard-curve TaqMan RT-qPCR using *in vitro*-transcribed HCV RNA, TaqMan Fast Virus 1-Step Master Mix (Invitrogen), and HCV-specific TaqMan primers (Invitrogen #4331182). Copy numbers were normalized to input volume and RNA concentration.

### Sanger RT-PCR

PCR was performed on cDNA with 2× CloneAmp HiFi PCR Premix (Takara). Cycling conditions were: 98°C/10 s, 55°C/5 s, 72°C/5 s for 35 cycles. Products were verified on 1.5% agarose gels and analyzed by Sanger sequencing (Azenta). Sequences were aligned to reference sequences and editing rates were quantified using EditR^66^.

### Native RIP-qPCR

Native RIP-qPCR was adapted from previously described methods^67^. For each condition, harvested cells were incubated in 400 µl native RIP buffer (NRB; 100 mM KCl, 5 mM MgCl_2_, 10 mM Tris-Cl pH 7.4, 0.5% Nonidet P-40 (NP-40; Thermo Scientific), protease inhibitor (Sigma-Aldrich), and 100 U/ml murine RNase inhibitor (NEB)). Following a 15 min incubation on ice, CaCl_2_ (5 mM final) and 45 U of DNase (NEB) were added and lysates were incubated at 25°C for 15 min with intermittent agitation (250 RPM, 1 min on/off). Lysates were cleared by centrifugation (16,000 × g, 15 min, 4°C), quantified by Bradford assay, and normalized to 3 mg/mL with NRB. For immunoprecipitation (IP), 300 µg lysate was incubated with either 10 µL rabbit V5-coupled magnetic beads or 2 µg rabbit IgG coupled to 10 µL Protein G Dynabeads for 1 h at room temperature in 500 µL NRB. Aliquots (20 µL and 50 µL) were reserved for protein and RNA inputs, respectively. Beads were washed 5× with 500 µL ice-cold NRB; 20% of beads were set aside for immunoblotting. Remaining beads and RNA inputs were digested with Proteinase K (2.4 U; NEB) at 50°C for 30 min (250 RPM, 1 min on/off) in 200 µL reactions (10 mM Tris-Cl pH 7.4, 100 mM NaCl, 12.5 mM ethylenediaminetetraacetic acid (EDTA), 0.1% sodium dodecyl sulfate (SDS), 1 µL GlycoBlue, 20 U RNasin Plus (Promega)), followed by phenol-chloroform extraction and ethanol precipitation. Following qPCR, enrichment was calculated as [% input V5 IP] / [% input IgG IP].

### Biotin-Labeled RNA Affinity

Biotin-labeled RNA affinity purification was adapted from established protocols^68,69^. Cell pellets were lysed 250 µL/plate ice-cold lysis buffer (10 mM Tris-Cl pH 7.5, 10 mM NaCl, 2 mM EDTA pH 8.0, 0.5% Triton X-100, protease inhibitor (Sigma-Aldrich), 0.5 mM DTT) and homogenized with 30 strokes in a pre-chilled type B dounce homogenizer. Nuclei were pelleted by centrifugation (max speed, 15 min, 4°C), and the cytoplasmic supernatant was adjusted to 5% glycerol and 150 mM KCl. Protein concentrations were determined by Bradford assay and normalized to 3 mg/mL. Lysates were pre-cleared with streptavidin agarose beads (100 µL/mg lysate, pre-washed in binding buffer) for 1 h at 4°C with rotation, then aliquoted (110 µL), snap-frozen, and stored at −80°C.

For RNA pulldowns, synthesized 38-nt biotinylated RNA probes (Horizon; 5’-biotin-GGGGCGUGUGGUCGGG[A/m⁶A]CUCGGCUUGGCUGCGCGUCCC-3’) were denatured at 95°C for 5 min, snap-cooled on ice, and immobilized on pre-washed streptavidin magnetic beads (250 pmol RNA per 5 µL beads) in 200 µL binding buffer (10 mM Tris-Cl pH 7.5, 1.5 mM MgCl₂, 150 mM KCl, 0.05% NP-40, protease inhibitor (Sigma-Aldrich), 0.5 mM DTT, 1 × 10⁵ U/mL murine RNase inhibitor (NEB-) for 2 h at 4°C with rotation. Beads were washed twice with binding buffer, resuspended in 400 µL binding buffer supplemented with RNase inhibitor, and incubated with 300 µg lysate for 30 min at 37°C with intermittent shaking (250 RPM, 1 min on/off). Beads were then washed 5× with binding buffer with brief vortexing at 50% power between washes, resuspended in 25 µL 1× SDS loading buffer, and denatured at 95°C for 5 min.

### Immunoblotting

Cell pellets were lysed in modified radioimmunoprecipitation assay buffer (TX-100-RIPA: 50 mM Tris pH 7.5, 150 mM NaCl, 5 mM EDTA, 0.1% SDS, 0.5% sodium deoxycholate, 1% Triton X-100) supplemented with protease inhibitor (Sigma-Aldrich). Lysates were cleared by centrifugation and quantified by Bradford assay (Bio-Rad). Samples were resolved by SDS-polyacrylamide gel electrophoresis (SDS-PAGE) and transferred to nitrocellulose membranes using a Trans-Blot Turbo system (Bio-Rad). Membranes were stained with Revert Total Protein Stain (LI-COR Biosciences), imaged on a LI-COR Odyssey FC, destained, and blocked with 3% bovine serum albumin (BSA) in Tris-buffered saline with 0.1% Tween-20 (TBS-T). Membranes were probed with primary antibodies, washed, incubated with species-specific horseradish peroxidase (HRP)-conjugated (Jackson ImmunoResearch; 1:5,000) or fluorescent (LI-COR; 1:5,000) secondary antibodies, and developed with Clarity Western ECL substrate (Bio-Rad). Imaging was performed on a LI-COR Odyssey FC.

Densitometry was performed with LI-COR Image Studio software and normalized using a two-step approach. For standard immunoblots, signals were first normalized to the corresponding total-protein stain for each lane, then normalized to the indicated within-experiment control condition (control set to 1) to yield relative values. For immunoprecipitation experiments, total-protein normalization was applied to input samples only. Signals in immunoprecipitated fractions were normalized to the corresponding total-protein–normalized input and then normalized to the designated control condition (control set to 1).

### Co-Immunoprecipitation (Co-IP) Assays

Cells (8 × 10⁵) were transfected with 1 µg of each plasmid using FuGENE 6 (Promega). At 24 h post-transfection, cells were harvested and lysed in NP-40 buffer (20 mM Tris pH 7.4, 100 mM NaCl, 0.5% NP-40) with protease inhibitor (Sigma-Aldrich). Cleared lysates (200 µg) were incubated with 20 µL anti-FLAG (Sigma-Aldrich) or anti-HA (Pierce/Thermo Fisher) magnetic beads for 1 h at 25°C with rotation. Beads were washed 3× with NP-40 buffer (8 min, rotating) and eluted in 2× SDS loading buffer with 5% β-mercaptoethanol (95°C, 5 min). Eluates and inputs were analyzed by immunoblotting.

### Recombinant Protein Purification

IFIT1 and IFIT3 were expressed individually in Escherichia coli Rosetta (DE3) cells (Novagen) grown in LB media to an OD₆₀₀ of 0.6 and induced with 0.5mM IPTG at 18 °C overnight. YTHDC1 was expressed in the same cells but grown in ZYM-5052 auto-induction media at 18 °C for 18 hours. The proteins were purified from clarified lysate using nickel affinity chromatography. For IFIT1, the SUMO tag was removed by overnight incubation with ULP1 protease followed by reverse Ni-NTA chromatography. Eluates from nickel affinity-chromatography were further purified by ion-exchange chromatography using a linear gradient from 100 mM to 1 M NaCl. Proteins were further purified by size-exclusion chromatography. The protein storage buffer for IFIT1 and IFIT3 contains 20 mM Tris-HCl, pH 7.5, 1 M NaCl, 10% (v/v) glycerol, and 2 mM DTT. The protein storage buffer for YTHDC1 contains 20 mM Bis-tris methane, pH 7, 1 M NaCl, 10% (v/v) glycerol and 5 mM DTT.

### Electrophoretic Mobility Shift Assay

38 nt RNA were synthesized ±m⁶A with 5’ triphosphate and 3’ biotin (Horizon; 5’-ppp-GGGGCGUGUGGUCGGG[A/m⁶A]CUCGGCUUGGCUGCGCGUCCC-biotin-3’). RNAs were 5’ radiolabeled using 0.4 U/µL T4 PNK in 1X PNK buffer (New England BioLabs) with 133 nM [γ-^32^P]ATP (Revvity). Radiolabeled RNAs were purified over Bio-Gel P-6 SEC beads (BioRad) equilibrated in 20 mM Tris-HCl, pH 7.5, 100 mM NaCl, and 2 mM MgCl_2_. RNA-protein complexes were formed by mixing the indicated concentrations of protein with 1 nM radiolabeled RNA substrate in 50 mM Tris-HCl, pH 7.5, 133mM NaCl, 10% glycerol, 5 mM DTT, and 2ng/µL of yeast tRNA (Roche) at 25 °C for 30 minutes, then resolved with a native 8% Tris-Borate-EDTA polyacrylamide gel. Gels were dried and imaged using a Typhoon FLA 9500 phosphorimager (GE Healthcare).

### Statistical Analysis

All experiments were performed with 3 biological replicates. Data are presented as mean ± SEM unless otherwise noted; for RNA-editing analyses, values represent mean editing. Graphs and statistical analyses were performed in GraphPad Prism (v10) unless stated otherwise. The following statistical tests were used: Empirical permutation testing with Benjamini–Hochberg FDR adjustment in Figures 1E-F, 2C-D (see also Methods Permutation Analyses of HyperTRIBE-seq); unpaired Student’s t test with single pooled variance and Holm-Šídák correction for multiple comparisons for Figure 2A; unpaired Student’s t test with individual variance and Holm-Šídák correction for multiple comparisons for Figure 2F, 3G-H, and 5D-E; paired two-tailed t test for Figure 3A; repeated measures one-way ANOVA with Geisser-Greenhouse and Dunnett’s correction for Figures 4C, 4I, and 5A-C. Significance thresholds are denoted as *p < 0.05, **p < 0.005, ***p < 0.0005, ****p < 0.00005.

## Notes

### Competing Interest Statement

The authors have declared no competing interest.

## REFERENCES

1. Schlee, M., and Hartmann, G. (2016). Discriminating self from non-self in nucleic acid sensing. Nat Rev Immunol 16, 566–580. 10.1038/nri.2016.78.

2. van Huizen, M., and Gack, M.U. (2025). The RIG-I-like receptor family of immune proteins. Molecular Cell 85, 3793–3806. 10.1016/j.molcel.2025.09.008.

3. Yang, E., and Li, M.M.H. (2020). All About the RNA: Interferon-Stimulated Genes That Interfere With Viral RNA Processes. Front Immunol 11, 605024. 10.3389/fimmu.2020.605024.

4. Schoggins, J.W. (2019). Interferon-Stimulated Genes: What Do They All Do? Annu Rev Virol 6, 567–584. 10.1146/annurev-virology-092818-015756.

5. Hur, S. (2019). Double-Stranded RNA Sensors and Modulators in Innate Immunity. Annu Rev Immunol 37, 349–375. 10.1146/annurev-immunol-042718-041356.

6. Diamond, M.S., and Farzan, M. (2013). The broad-spectrum antiviral functions of IFIT and IFITM proteins. Nat Rev Immunol 13, 46–57. 10.1038/nri3344.

7. Mears, H.V., and Sweeney, T.R. (2018). Better together: the role of IFIT protein-protein interactions in the antiviral response. J Gen Virol 99, 1463–1477. 10.1099/jgv.0.001149.

8. Shaw, A.E., Hughes, J., Gu, Q., Behdenna, A., Singer, J.B., Dennis, T., Orton, R.J., Varela, M., Gifford, R.J., Wilson, S.J., and Palmarini, M. (2017). Fundamental properties of the mammalian innate immune system revealed by multispecies comparison of type I interferon responses. PLoS Biol 15, e2004086. 10.1371/journal.pbio.2004086.

9. Zhou, X., Michal, J.J., Zhang, L., Ding, B., Lunney, J.K., Liu, B., and Jiang, Z. (2013). Interferon induced IFIT family genes in host antiviral defense. Int J Biol Sci 9, 200–208. 10.7150/ijbs.5613.

10. Liu, Y., Zhang, Y.B., Liu, T.K., and Gui, J.F. (2013). Lineage-specific expansion of IFIT gene family: an insight into coevolution with IFN gene family. PLoS One 8, e66859. 10.1371/journal.pone.0066859.

11. Sullivan, O.M., Nesbitt, D.J., Schaack, G.A., Feltman, E.M., Nipper, T., Kongsomros, S., Reed, S.G., Nelson, S.L., King, C.R., Shishkova, E., et al. (2025). IFIT3 RNA-binding activity promotes influenza A virus infection and translation efficiency. Journal of Virology 99, e00286–00225. doi:10.1128/jvi.00286-25.

12. Glasner, D.R., Todd, C., Cook, B., D’Urso, A., Khosla, S., Estrada, E., Wagner, J.D., Bartels, M.D., Hung, C.T., Ford, P., et al. (2025). The IFIT2-IFIT3 antiviral complex targets short 5’ untranslated regions on viral mRNAs for translation inhibition. Nat Microbiol. 10.1038/s41564-025-02138-w.

13. Clayton, E., Atasoy, M.O., Naggar, R.F.E., Franco, A.C., Rohaim, M.A., and Munir, M. (2024). Interferon-induced protein with tetratricopeptide repeats 5 of black fruit bat (Pteropus alecto) displays a broad inhibition of RNA viruses. Front Immunol 15, 1284056. 10.3389/fimmu.2024.1284056.

14. Tran, V., Ledwith, M.P., Thamamongood, T., Higgins, C.A., Tripathi, S., Chang, M.W., Benner, C., García-Sastre, A., Schwemmle, M., Boon, A.C.M., et al. (2020). Influenza virus repurposes the antiviral protein IFIT2 to promote translation of viral mRNAs. Nature Microbiology. 10.1038/s41564-020-0778-x.

15. Ishida, Y., Kakuni, M., Bang, B.R., Sugahara, G., Lau, D.T., Tateno-Mukaidani, C., Li, M., Gale, M., Jr., and Saito, T. (2019). Hepatic IFN-Induced Protein with Tetratricopeptide Repeats Regulation of HCV Infection. J Interferon Cytokine Res 39, 133–146. 10.1089/jir.2018.0103.

16. Johnson, B., VanBlargan, L.A., Xu, W., White, J.P., Shan, C., Shi, P.Y., Zhang, R., Adhikari, J., Gross, M.L., Leung, D.W., et al. (2018). Human IFIT3 Modulates IFIT1 RNA Binding Specificity and Protein Stability. Immunity 48, 487–499.e485. 10.1016/j.immuni.2018.01.014.

17. Fleith, R.C., Mears, H.V., Leong, X.Y., Sanford, T.J., Emmott, E., Graham, S.C., Mansur, D.S., and Sweeney, T.R. (2018). IFIT3 and IFIT2/3 promote IFIT1-mediated translation inhibition by enhancing binding to non-self RNA. Nucleic Acids Res 46, 5269–5285. 10.1093/nar/gky191.

18. Yang, Z., Liang, H., Zhou, Q., Li, Y., Chen, H., Ye, W., Chen, D., Fleming, J., Shu, H., and Liu, Y. (2012). Crystal structure of ISG54 reveals a novel RNA binding structure and potential functional mechanisms. Cell Res 22, 1328–1338. 10.1038/cr.2012.111.

19. Poddar, D., Sharma, N., Ogino, T., Qi, X., Kessler, P.M., Mendries, H., Dutta, R., and Sen, G.C. (2024). The interferon-induced protein, IFIT2, requires RNA-binding activity and neuronal expression to protect mice from intranasal vesicular stomatitis virus infection. mBio 15, e0056824. 10.1128/mbio.00568-24.

20. Miedziak, B., Dobiezynska, A., Darzynkiewicz, Z.M., Bartkowska, J., Miszkiewicz, J., Kowalska, J., Warminski, M., Tyras, M., Trylska, J., Jemielity, J., et al. (2020). Kinetic analysis of IFIT1 and IFIT5 interactions with different native and engineered RNAs and its consequences for designing mRNA-based therapeutics. RNA 26, 58–68. 10.1261/rna.073304.119.

21. Katibah, G.E., Lee, H.J., Huizar, J.P., Vogan, J.M., Alber, T., and Collins, K. (2013). tRNA binding, structure, and localization of the human interferon-induced protein IFIT5. Mol Cell 49, 743–750. 10.1016/j.molcel.2012.12.015.

22. Katibah, G.E., Qin, Y., Sidote, D.J., Yao, J., Lambowitz, A.M., and Collins, K. (2014). Broad and adaptable RNA structure recognition by the human interferon-induced tetratricopeptide repeat protein IFIT5. Proc Natl Acad Sci U S A 111, 12025–12030. 10.1073/pnas.1412842111.

23. Feng, F., Yuan, L., Wang, Y.E., Crowley, C., Lv, Z., Li, J., Liu, Y., Cheng, G., Zeng, S., and Liang, H. (2013). Crystal structure and nucleotide selectivity of human IFIT5/ISG58. Cell Res 23, 1055–1058. 10.1038/cr.2013.80.

24. Metz, P., Dazert, E., Ruggieri, A., Mazur, J., Kaderali, L., Kaul, A., Zeuge, U., Windisch, M.P., Trippler, M., Lohmann, V., et al. (2012). Identification of type I and type II interferon-induced effectors controlling hepatitis C virus replication. Hepatology 56, 2082–2093. 10.1002/hep.25908.

25. Rahman, R., Xu, W., Jin, H., and Rosbash, M. (2018). Identification of RNA-binding protein targets with HyperTRIBE. Nat Protoc 13, 1829–1849. 10.1038/s41596-018-0020-y.

26. Xu, W., Rahman, R., and Rosbash, M. (2018). Mechanistic implications of enhanced editing by a HyperTRIBE RNA-binding protein. Rna 24, 173–182. 10.1261/rna.064691.117.

27. Kuttan, A., and Bass, B.L. (2012). Mechanistic insights into editing-site specificity of ADARs. Proc Natl Acad Sci U S A 109, E3295–3304. 10.1073/pnas.1212548109.

28. Abruzzi, K.C., Ratner, C., and Rosbash, M. (2023). Comparison of TRIBE and STAMP for identifying targets of RNA binding proteins in human and Drosophila cells. Rna 29, 1230–1242. 10.1261/rna.079608.123.

29. Zhu, H., Yin, X., Holley, C.L., and Meyer, K.D. (2022). Improved Methods for Deamination-Based m(6)A Detection. Front Cell Dev Biol 10, 888279. 10.3389/fcell.2022.888279.

30. Tegowski, M., Flamand, M.N., and Meyer, K.D. (2022). scDART-seq reveals distinct m(6)A signatures and mRNA methylation heterogeneity in single cells. Mol Cell 82, 868–878 e810. 10.1016/j.molcel.2021.12.038.

31. Flamand, M.N., and Meyer, K.D. (2022). m6A and YTHDF proteins contribute to the localization of select neuronal mRNAs. Nucleic Acids Research 50, 4464–4483. 10.1093/nar/gkac251.

32. Luo, E.C., Nathanson, J.L., Tan, F.E., Schwartz, J.L., Schmok, J.C., Shankar, A., Markmiller, S., Yee, B.A., Sathe, S., Pratt, G.A., et al. (2020). Large-scale tethered function assays identify factors that regulate mRNA stability and translation. Nat Struct Mol Biol. 10.1038/s41594-020-0477-6.

33. Zhao, W., Zhang, S., Zhu, Y., Xi, X., Bao, P., Ma, Z., Kapral, Thomas H., Chen, S., Zagrovic, B., Yang, Yucheng T., and Lu, Zhi J. (2021). POSTAR3: an updated platform for exploring post-transcriptional regulation coordinated by RNA-binding proteins. Nucleic Acids Research 50, D287–D294. 10.1093/nar/gkab702.

34. Mitschka, S., and Mayr, C. (2022). Context-specific regulation and function of mRNA alternative polyadenylation. Nat Rev Mol Cell Biol 23, 779–796. 10.1038/s41580-022-00507-5.

35. Huang da, W., Sherman, B.T., and Lempicki, R.A. (2009). Systematic and integrative analysis of large gene lists using DAVID bioinformatics resources. Nat Protoc 4, 44–57. 10.1038/nprot.2008.211.

36. Dominissini, D., Moshitch-Moshkovitz, S., Schwartz, S., Salmon-Divon, M., Ungar, L., Osenberg, S., Cesarkas, K., Jacob-Hirsch, J., Amariglio, N., Kupiec, M., et al. (2012). Topology of the human and mouse m6A RNA methylomes revealed by m6A-seq. Nature 485, 201–206. 10.1038/nature11112.

37. Cai, Z., Xu, H., Bai, G., Hu, H., Wang, D., Li, H., and Wang, Z. (2022). ELAVL1 promotes prostate cancer progression by interacting with other m6A regulators. Front Oncol 12, 939784. 10.3389/fonc.2022.939784.

38. McMillan, M., Gomez, N., Hsieh, C., Bekier, M., Li, X., Miguez, R., Tank, E.M.H., and Barmada, S.J. (2023). RNA methylation influences TDP43 binding and disease pathogenesis in models of amyotrophic lateral sclerosis and frontotemporal dementia. Mol Cell. 10.1016/j.molcel.2022.12.019.

39. Mehravar, M., Kumar, Y., Olshansky, M., Dakle, P., Bullen, M., Pandey, V.K., Bansal, D., Dent, C., Hathiwala, D., Zhang, Z., et al. (2022). MOV10 facilitates messenger RNA decay in an N6-methyladenosine (m^6^A) dependent manner to maintain the mouse embryonic stem cells state. bioRxiv, 2021.2008.2011.456030. 10.1101/2021.08.11.456030.

40. Edens, B.M., Vissers, C., Su, J., Arumugam, S., Xu, Z., Shi, H., Miller, N., Rojas Ringeling, F., Ming, G.L., He, C., et al. (2019). FMRP Modulates Neural Differentiation through m(6)A-Dependent mRNA Nuclear Export. Cell Rep 28, 845–854 e845. 10.1016/j.celrep.2019.06.072.

41. Edupuganti, R.R., Geiger, S., Lindeboom, R.G.H., Shi, H., Hsu, P.J., Lu, Z., Wang, S.Y., Baltissen, M.P.A., Jansen, P., Rossa, M., et al. (2017). N(6)-methyladenosine (m(6)A) recruits and repels proteins to regulate mRNA homeostasis. Nat Struct Mol Biol 24, 870–878. 10.1038/nsmb.3462.

42. Sacco, M.T., Bland, K.M., and Horner, S.M. (2022). WTAP Targets the METTL3 m(6)A-Methyltransferase Complex to Cytoplasmic Hepatitis C Virus RNA to Regulate Infection. J Virol 96, e0099722. 10.1128/jvi.00997-22.

43. Gokhale, N.S., McIntyre, A.B.R., McFadden, M.J., Roder, A.E., Kennedy, E.M., Gandara, J.A., Hopcraft, S.E., Quicke, K.M., Vazquez, C., Willer, J., et al. (2016). N6-Methyladenosine in Flaviviridae Viral RNA Genomes Regulates Infection. Cell Host Microbe 20, 654–665. 10.1016/j.chom.2016.09.015.

44. Yankova, E., Blackaby, W., Albertella, M., Rak, J., De Braekeleer, E., Tsagkogeorga, G., Pilka, E.S., Aspris, D., Leggate, D., Hendrick, A.G., et al. (2021). Small-molecule inhibition of METTL3 as a strategy against myeloid leukaemia. Nature 593, 597–601. 10.1038/s41586-021-03536-w.

45. Wang, X., Lu, Z., Gomez, A., Hon, G.C., Yue, Y., Han, D., Fu, Y., Parisien, M., Dai, Q., Jia, G., et al. (2014). N6-methyladenosine-dependent regulation of messenger RNA stability. Nature 505, 117–120. 10.1038/nature12730.

46. Gokhale, N.S., McIntyre, A.B.R., Mattocks, M.D., Holley, C.L., Lazear, H.M., Mason, C.E., and Horner, S.M. (2020). Altered m(6)A Modification of Specific Cellular Transcripts Affects Flaviviridae Infection. Mol Cell 77, 542–555 e548. 10.1016/j.molcel.2019.11.007.

47. Wan, H., Adams, R.L., Lindenbach, B.D., and Pyle, A.M. (2022). The In Vivo and In Vitro Architecture of the Hepatitis C Virus RNA Genome Uncovers Functional RNA Secondary and Tertiary Structures. J Virol 96, e0194621. 10.1128/jvi.01946-21.

48. Abramson, J., Adler, J., Dunger, J., Evans, R., Green, T., Pritzel, A., Ronneberger, O., Willmore, L., Ballard, A.J., Bambrick, J., et al. (2024). Accurate structure prediction of biomolecular interactions with AlphaFold 3. Nature 630, 493–500. 10.1038/s41586-024-07487-w.

49. Abbas, Y.M., Laudenbach, B.T., Martinez-Montero, S., Cencic, R., Habjan, M., Pichlmair, A., Damha, M.J., Pelletier, J., and Nagar, B. (2017). Structure of human IFIT1 with capped RNA reveals adaptable mRNA binding and mechanisms for sensing N1 and N2 ribose 2’-O methylations. Proc Natl Acad Sci U S A 114, E2106–E2115. 10.1073/pnas.1612444114.

50. Geng, J., Chrabaszczewska, M., Kurpiejewski, K., Stankiewicz-Drogon, A., Jankowska-Anyszka, M., Darzynkiewicz, E., and Grzela, R. (2024). Cap-related modifications of RNA regulate binding to IFIT proteins. RNA 30, 1292–1305. 10.1261/rna.080011.124.

51. Lindenbach, B.D., and Rice, C.M. (2013). The ins and outs of hepatitis C virus entry and assembly. Nat Rev Microbiol 11, 688–700. 10.1038/nrmicro3098.

52. Baquero-Perez, B., Geers, D., and Diez, J. (2021). From A to m(6)A: The Emerging Viral Epitranscriptome. Viruses 13. 10.3390/v13061049.

53. Iselin, L., Demyanenko, Y., Palmalux, N., Buh, A.E., Kamel, W., Simmonds, P., Mohammed, S., and Castello, A. (2025). RNA binding regulation is a new dimension in the type I IFN response. bioRxiv, 2025.2005.2030.656608. 10.1101/2025.05.30.656608.

54. Garcia-Moreno, M., Noerenberg, M., Ni, S., Järvelin, A.I., González-Almela, E., Lenz, C.E., Bach-Pages, M., Cox, V., Avolio, R., Davis, T., et al. (2019). System-wide Profiling of RNA-Binding Proteins Uncovers Key Regulators of Virus Infection. Mol Cell 74, 196–211.e111. 10.1016/j.molcel.2019.01.017.

55. Liepelt, A., Naarmann-de Vries, I.S., Simons, N., Eichelbaum, K., Föhr, S., Archer, S.K., Castello, A., Usadel, B., Krijgsveld, J., Preiss, T., et al. (2016). Identification of RNA-binding Proteins in Macrophages by Interactome Capture. Mol Cell Proteomics 15, 2699–2714. 10.1074/mcp.M115.056564.

56. Lu, M., Zhang, Z., Xue, M., Zhao, B.S., Harder, O., Li, A., Liang, X., Gao, T.Z., Xu, Y., Zhou, J., et al. (2020). N(6)-methyladenosine modification enables viral RNA to escape recognition by RNA sensor RIG-I. Nat Microbiol 5, 584–598. 10.1038/s41564-019-0653-9.

57. Kim, G.W., Imam, H., Khan, M., and Siddiqui, A. (2020). N (6)-Methyladenosine modification of hepatitis B and C viral RNAs attenuates host innate immunity via RIG-I signaling. J Biol Chem 295, 13123–13133. 10.1074/jbc.RA120.014260.

58. Aligeti, M., Roder, A., and Horner, S.M. (2015). Cooperation between the Hepatitis C Virus p7 and NS5B Proteins Enhances Virion Infectivity. J Virol 89, 11523–11533. 10.1128/JVI.01185-15.

59. Sherwood, A.V., Rivera-Rangel, L.R., Ryberg, L.A., Larsen, H.S., Anker, K.M., Costa, R., Vagbo, C.B., Jakljevic, E., Pham, L.V., Fernandez-Antunez, C., et al. (2023). Hepatitis C virus RNA is 5’-capped with flavin adenine dinucleotide. Nature. 10.1038/s41586-023-06301-3.

60. Di Tommaso, P., Chatzou, M., Floden, E.W., Barja, P.P., Palumbo, E., and Notredame, C. (2017). Nextflow enables reproducible computational workflows. Nat Biotechnol 35, 316–319. 10.1038/nbt.3820.

61. Ewels, P.A., Peltzer, A., Fillinger, S., Patel, H., Alneberg, J., Wilm, A., Garcia, M.U., Di Tommaso, P., and Nahnsen, S. (2020). The nf-core framework for community-curated bioinformatics pipelines. Nat Biotechnol 38, 276–278. 10.1038/s41587-020-0439-x.

62. Gruning, B., Dale, R., Sjodin, A., Chapman, B.A., Rowe, J., Tomkins-Tinch, C.H., Valieris, R., Koster, J., and Bioconda, T. (2018). Bioconda: sustainable and comprehensive software distribution for the life sciences. Nat Methods 15, 475–476. 10.1038/s41592-018-0046-7.

63. da Veiga Leprevost, F., Gruning, B.A., Alves Aflitos, S., Rost, H.L., Uszkoreit, J., Barsnes, H., Vaudel, M., Moreno, P., Gatto, L., Weber, J., et al. (2017). BioContainers: an open-source and community-driven framework for software standardization. Bioinformatics 33, 2580–2582. 10.1093/bioinformatics/btx192.

64. Sherman, B.T., Hao, M., Qiu, J., Jiao, X., Baseler, M.W., Lane, H.C., Imamichi, T., and Chang, W. (2022). DAVID: a web server for functional enrichment analysis and functional annotation of gene lists (2021 update). Nucleic Acids Res 50, W216–w221. 10.1093/nar/gkac194.

65. Quinlan, A.R., and Hall, I.M. (2010). BEDTools: a flexible suite of utilities for comparing genomic features. Bioinformatics 26, 841–842. 10.1093/bioinformatics/btq033.

66. Kluesner, M.G., Nedveck, D.A., Lahr, W.S., Garbe, J.R., Abrahante, J.E., Webber, B.R., and Moriarity, B.S. (2018). EditR: A Method to Quantify Base Editing from Sanger Sequencing. The CRISPR Journal 1, 239–250. 10.1089/crispr.2018.0014.

67. Conrad, N.K. (2008). Chapter 15. Co-immunoprecipitation techniques for assessing RNA-protein interactions in vivo. Methods Enzymol 449, 317–342. 10.1016/S0076-6879(08)02415-4.

68. Choi, S.H., Flamand, M.N., Liu, B., Zhu, H., Hu, M., Wang, M., Sewell, J., Holley, C.L., Al-Hashimi, H.M., and Meyer, K.D. (2022). RBM45 is an m(6)A-binding protein that affects neuronal differentiation and the splicing of a subset of mRNAs. Cell Rep 40, 111293. 10.1016/j.celrep.2022.111293.

69. Thompson, M.G., Munoz-Moreno, R., Bhat, P., Roytenberg, R., Lindberg, J., Gazzara, M.R., Mallory, M.J., Zhang, K., Garcia-Sastre, A., Fontoura, B.M.A., and Lynch, K.W. (2018). Co-regulatory activity of hnRNP K and NS1-BP in influenza and human mRNA splicing. Nat Commun 9, 2407. 10.1038/s41467-018-04779-4.

